# The Neural Consequences of Attentional Prioritization of Internal Representations in Visual Working Memory

**DOI:** 10.1101/606665

**Authors:** Muhammet I. Sahan, Andrew D. Sheldon, Bradley R. Postle

**Affiliations:** Department of Psychiatry, University of Wisconsin, Madison, WI, USA; Neuroscience Training Program, University of Wisconsin, Madison, WI, USA; Medical Scientist Training Program, University of Wisconsin, Madison, WI, USA; Department of Experimental Psychology, Ghent University, Ghent, Belgium; Department of Psychology, University of Wisconsin, Madison, WI, USA

**Author notes:** These authors contributed equally to this work (Muhammet I. Sahan, Andrew D. Sheldon).

## Abstract

Although humans can hold multiple items in mind simultaneously, the contents of working memory (WM) can be selectively prioritized to effectively guide behavior in response to rapidly changing exigencies in the environment. Neural evidence for this is seen in studies of dual serial retrocuing of two items held concurrently in visual WM, in which evidence in occipital cortex for the active neural representation of the cued item increases, and evidence for the uncued item decreases, often to levels indistinguishable from empirical baseline. Although this pattern is reminiscent of the effects of selective attention on visual perception, the extent to which more subtle principles of visual attention may also apply to visual working memory remains uncertain. In the present study we explored whether the well-characterized “same-object” benefit in visual target detection, attributed to object-based attention (e.g., Duncan, 1984; Egly, Driver, & Rafal, 1994), may also be observed for information held in visual WM. fMRI data were collected while human subjects (male and female) performed a multi-step serial retrocuing task in which they first viewed two two-dimensional sample stimuli comprised of colored moving dots. After stimulus offset, an initial *relevance cue* then indicated whether both dimensions of only the first or only the second object, or only the color or only the direction-of-motion of both objects, would be relevant for the remainder of the trial, which then proceeded with the standard dual serial retrocuing procedure. Thus, on “object-relevant” trials, the ensuing *priority cues* prompted the selection of one from among two features (“color” or “direction”) bound to the same object, whereas on “feature-relevant” trials the *priority cues* prompted the selection of one from among two features each belonging to a different object. Results of analyses with multivariate inverted encoding models (IEM) revealed a same-object benefit on object-relevant trials: Whereas, on feature-relevant trials, the first *priority cue* triggered a strengthening of the neural representation of the cued feature and a concomitant weakening-to-baseline of the uncued feature; on object-relevant trials the cued item remained active but did not increase in strength, and the uncued item weakened, but remained significantly elevated throughout the delay period. Of additional interest, on both types of trials the second *priority cue* prompted an active recoding of the uncued item into a different neural representation, perhaps to minimize its ability to interfere with recall of the cued item. Finally, although stimulus-specific representation in parietal and frontal cortex was weak and uneven, these regions closely tracked the higher-order information of which stimulus category was relevant for behavior at all points during the trial, indicating an important role in controlling the prioritization of information in visual working memory.

## Introduction

Working memory (WM) is a cognitive function that enables the mental retention of information, in the absence of sustained input from the physical world, its manipulation, and its use for guiding behavior. Current accounts of sensory WM hold that it relies on attentional mechanisms also involved in the prioritization of information perceived in the environment (D’Esposito & Postle, 2015; Gazzaley & Nobre, 2012; Kiyonaga & Egner, 2013; Oberauer & Hein, 2012). Consistent with this view is that fact that instructing an individual to prioritize a subset of information being held in WM with a retrodictive cue (hereafter, “retrocue”) improves its subsequent recall at the expense of uncued information, in a manner comparable to the effects on visual perception of prospectively cuing a location or a feature in an impending visual scene (Griffin & Nobre, 2003; Lepsien & Nobre, 2007; Pertzov, Bays, Joseph, & Husain, 2013; Zokaei, Manohar, Husain, & Feredoes, 2013; Sahan, Verguts, Boehler, Pourtois & Fias, 2015). The retrocuing technique has also been used to test theories about the capacity and the temporal dynamics of the putative focus of attention (or “region of direct access”) in state-based models of WM (as reviewed in Larocque, Lewis-Peacock, & Postle (2014)). In one series of studies we have used a multistep delayed serial retrocuing (DSR) procedure in which subjects are first presented with two sample items to hold in WM, then, after a retention interval, an initial retrocue indicates which of the two will be tested by the impending memory probe. Because subjects know, however, that this probe will be followed by a second retrocue and a second probe, and because they know there is an equal probability that the second retrocue will prioritize either sample item, they cannot forget the item not cued by the initial retrocue. This creates a portion of the trial in which two items are being held in WM, but only one is a “prioritized memory item” [PMI]). Initially, in studies with fMRI (Larocque, Riggall, Emrich, & Postle, 2017; Lewis-Peacock & Postle, 2012) and with EEG (LaRocque, Lewis-Peacock, Drysdale, Oberauer, & Postle, 2013; Rose et al., 2016), multivariate pattern analyses (MVPA) showed that decoding the neural representation of the PMI improved following the initial retrocue, whereas decoding the neural representation of the initially uncued (and, therefore, “unprioritized”) memory item (UMI) dropped to baseline levels. More recently, there have been reports of multivariate evidence for the UMI, with an item represented in a different region (e.g., Christophel, Iamshchinina, Yan, Allefeld, & Haynes, 2018) or in a different neural code (e.g., van Loon, Olmos-solis, Fahrenfort, & Olivers, 2018) when a UMI relative to when a PMI. The present study was designed to use a multivariate inverted encoding modelling (IEM) approach, which offers advantages over multivariate decoding approaches (Serences & Saproo, 2012), to address several questions that arise from these observations: 1) is one effect of selection to increase the strength of the neural representation of the PMI? 2) does the degradation of MVPA decodability of the UMI truly correspond to a weakening of its neural representation? 3) regardless of the answer to (2), is the retrocuing effect on the neural representation of the UMI sensitive to its status as a discrete object or as a feature in a multidimensional object? The answers to these questions will have implications for broader questions, such as whether principles of object- and feature-based attention also apply to WM, and whether a complex object is represented in WM as more than the set of features that define it.

Well-established principles of visual attention, such biased-competition (Desimone & Duncan, 1995) and divisive normalization (Carandini & Heeger, 2011), have the potential to account the finding that retrocuing leads to a weakening of the UMI. That is, prioritization of one among multiple mnemonic representations could be achieved via top-down signals from frontoparietal systems that are important for the endogenous control of attention (e.g., Nelissen, Stokes, Nobre, & Rushworth (2013)). Importantly, the dynamics of biased competition have been demonstrated, with MVPA, to influence the population-level representation of objects in a manner that would be predicted from single-neuron studies, both in analyses of extracellular recordings from neurons in monkey ventral temporal cortex (Zhang, Meyers, Bichot, Serre, Poggio, & Desimone (2011)) and in unpublished fMRI data from humans performing a selective attention task (Sheldon, Saad, Sahan, Meyering & Postle, 2017). Indeed, it is the effects suggesting the operation of object-based attention, in a different previous fMRI study of WM, that motivate the present experiment.

In a DSR study by Lewis-Peacock, Drysdale and Postle (2014), subjects first viewed an image of a real-world object (e.g., a baseball), and were then cued as to what dimension of that stimulus would be interrogated by a memory probe: its silhouette outline; the phonology of its name; or the semantic category to which it belonged. Although the performance of classifiers trained independently to discriminate visual from phonological from semantic processing strengthened and weakened in a manner congruent with the cues, unlike in studies that required WM for two discrete objects, decoding for the uncued stimulus features did not drop to baseline levels. One possible account of this observation was that we were observing a neural correlate of an analogue of the “same-object” benefit that is seen in visual selective detection (Driver, 2001; Duncan, 1984; Vecera, Behrmann, & McGoldrick, 2000). That is, perhaps the uncued stimulus dimensions retained some level of activity because they were an inherent part of the same object from which a different stimulus dimension had been selected, and attention therefore spread to all components of the selected object. Limitations of that study’s design, however, precluded a strong test of this possibility.

The question of whether multidimensional objects are represented as bound objects in visual WM remains contentious in WM research. On one hand, there are several studies suggesting that objects defined by a conjunction of two or more features can be maintained just as well as can single-feature objects, suggesting that the elementary units of visual WM are integrated objects (e.g., Luck & Vogel, 1997; Luria & Vogel, 2011; Woodman & Vogel, 2008). Others have argued, however, that the elementary units of visual WM are the features that make up complex visual objects, and that the various features of an object are simultaneously stored in dimension-specific channels (e.g., Bays, Wu, & Husain, 2011; Wheeler & Treisman, 2002). Furthermore, according to these feature-based accounts, feature binding in WM only occurs when attention is exerted over the to-be-bound features. For instance, Wheeler and Treisman (2002) showed that same-object benefits were observed in visual WM only when subjects were not holding competing multi-feature objects in WM, presumably because these would disrupt sustained attentional control. The design of the present study may also help to address this debate.

To address more directly whether principles of object-based attention can be observed during visual WM, we designed the present study to compare the neural effects of selecting one feature from among two 1-dimensional objects being held concurrently in WM, versus those of selecting one feature from a single two-dimensional compound object being held in WM. Additionally, for the present study we adopted an analytic method that would allow us quantify the effects of selection on the strength of WM representations. A limitation of the MVPA decoding approach, such as what we have used in many previous DSR studies, is that it does not provide a direct measure of neural representations. Thus, for example, although one can observe systematic changes in MVPA performance with the manipulation of, say, the number of items being held in WM (e.g., Emrich, Riggall, Larocque, & Postle (2013)), the interpretation of such a finding is equivocal. Although it could be the strength of the neural representations of items that declines with increasing load, there are equally plausible alternative explanations: Perhaps increasing load changes the level of stimulus-nonspecific noise that nonetheless influences the performance of the decoder; or perhaps increasing load changes the nature of the neural code, but not of the amplitude per se, of stimulus representations. Inverted *encoding* modeling (IEM), in contrast, entails the fitting of data to one or more a priori models that specify the mapping from multiple sensor-level signals into a hypothesized population-level representation. This affords the quantification of different parameters of the model fit, such that one can estimate, in our case, the extent to which selection might strengthen or weaken neural representations in different experimental conditions. Additionally, testing the same data with different models can provide evidence for changes to the neural code.

## Methods

### Subjects

Ten neurologically healthy students from the University of Wisconsin–Madison (3 females, 18-30 years, *M* = 22, all right-handed) participated in three 2-hour scanning sessions. One subject was excluded from the analyses due to excessive head movement. Another subject was an author of this study (A.D.S.). All subjects had normal or corrected-to-normal vision and reported having normal color vision. The research complied to the guidelines of the University of Wisconsin–Madison’s Health Sciences Institutional Review Board, and all subjects gave written informed consent.

### Design

The experiment comprised a 1-item delayed-recall (a.k.a. “delayed-estimation”) task and a multiple serial retrocuing task (MSR; Figure 1). The purpose of the delayed-recall task was twofold: to serve as a localizer that was independent of the MSR task; and to train feature dimension-specific IEMs for testing on the MSR data. Importantly, fits to such an “independent” IEM can be used for quantitative comparisons between experimental conditions of interest (e.g., the effect of priority status on the neural representation of direction-of-motion in the MSR task). The delayed-recall task began with the presentation of a sample stimulus (2 sec), either a patch of uniformly colored static dots whose color varied from trial to trial, or a patch of grey dots moving with 100% coherence in a direction that varied from trial to trial. Data from this task was used to train IEMs to learn the neural bases of perceiving and remembering “color” and “direction of motion”. We note that it is possible that, when moving dots are presented at high contrast within a circular aperture, as is the case here, it is possible that the signals that support direction-of-motion decoding may not (only) correspond to motion processing, but, perhaps (also) to other factors, such as the transients generated by appearance/disappearance of individual dots at the opaque boundary of the aperture (e.g., an aperture-inward bias, Wang, Merriam, Freeman, & Heeger, (2014)). Importantly, the possible ambiguity about the precise computations underlying the signals generated by in this condition are not problematic for interpreting our results, because our interest is not in the neural bases of motion perception, per se, but rather in the neural bases of attention to either of two visual features that had the subjective properties (for the subject) of being categorically different – one reproducible on a color bar, the other with a radial dial -- and the objective properties (for the experimenters) of being varied along orthogonal dimensions – values in color space for static dots or direction of motion for color-invariant gray dots. For expository parsimony, from this point onward we will refer to these feature dimensions as “color” and “direction.”

**Figure 1.**
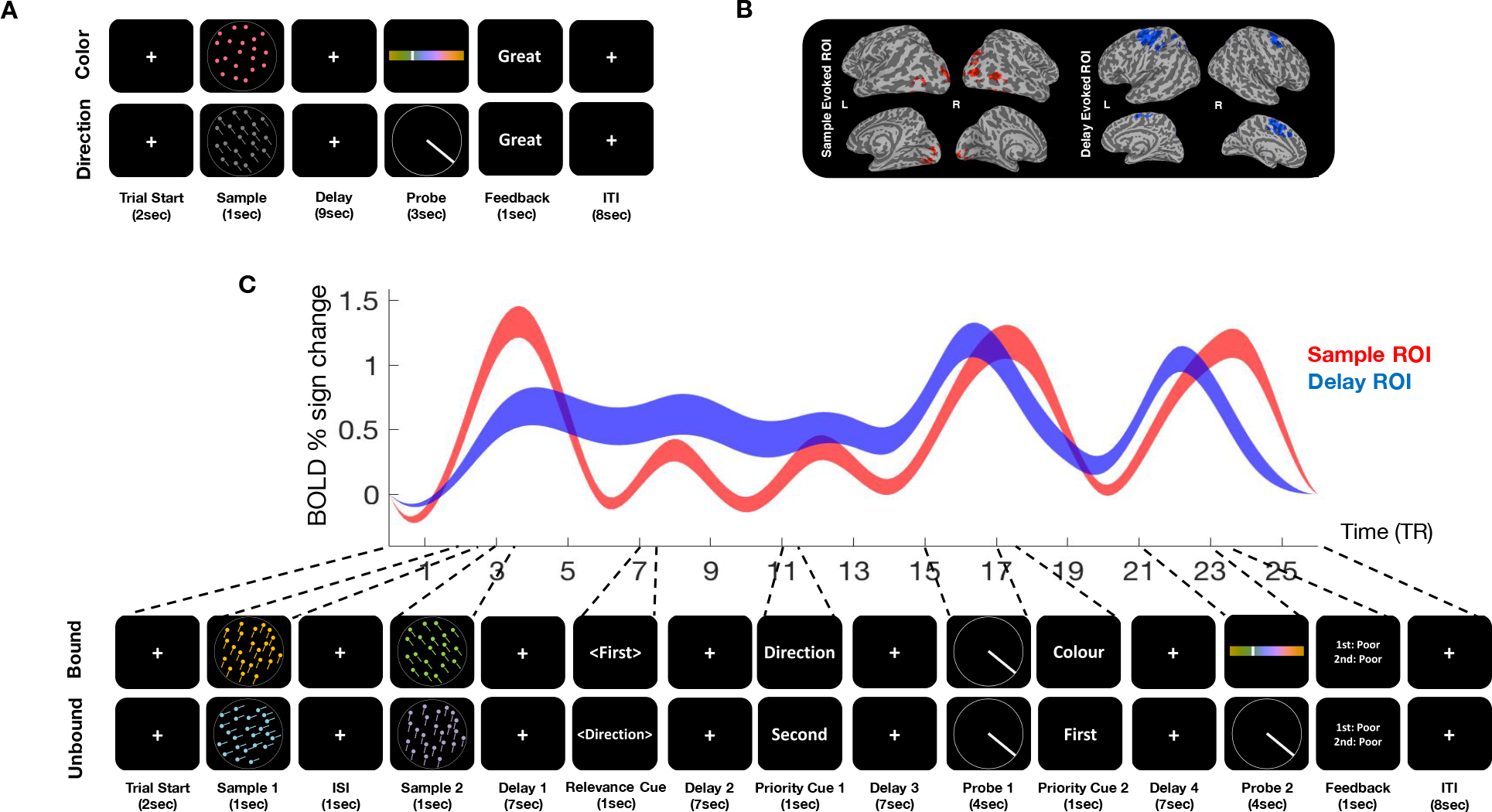
(A) Illustration of a trial from each condition of the 1-item delayed recall task. (B) Sample-evoked feature-nonselective VOT ROI, and Delay-evoked feature-nonselective frontal ROI, from a representative subject. (C) Illustration of a trial from the *Bound* and a trial from *Unbound* conditions of the MSR task, together with group-, condition-, and trial-averaged BOLD signal from the ROIs illustrated in B. The width of the traces denotes the standard error of the mean.

**Figure 2.**
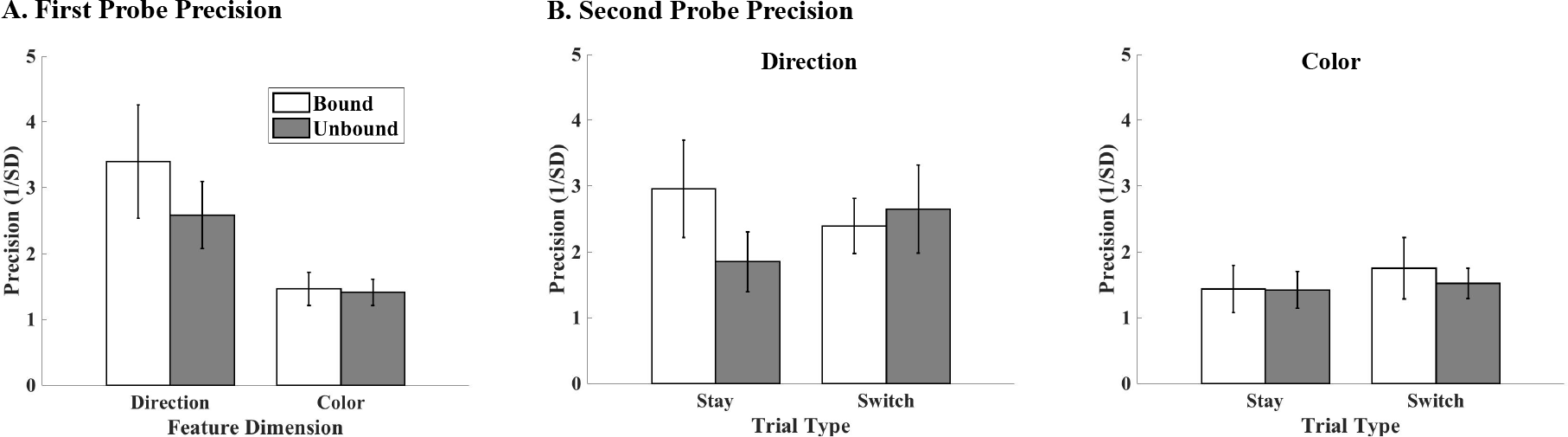
Model-free behavioral results. (A) *Probe 1*: Recall was less precise for color than for direction trials, but insensitive to binding status. (B) *Probe 2*: Recall was less precise for color than for direction trials, and insensitive to binding condition and to “stay”/“switch” status of *priority cue 2*.

Each trial of the task of primary experimental interest, the MSR task, began with the serial presentation of two 2-dimensional stimuli, each a patch of coherently moving colored dots. Next, a “relevance cue” designated what information from the two sample stimuli would be relevant for that trial: the color and direction of either the first or the second sample; or the color of both samples or the direction of both samples. Thus, a *relevance cue* indicating “<First>” or “<Second>” would designate a trial that would require selection of a feature bound to a 2-D object, whereas a *relevance cue* indicating “<Color>” or “<Direction>” would designate a trial that would require selection of the relevant feature of one from among two objects. Following the *relevance cue*, each trial proceeded in the same way as many previous D(ual)SR tasks: a first “prioritization cue” indicated which of the two relevant features would be tested by the first recall probe, then a second *prioritization cue* indicated, with a probability of .5, which of the same two relevant features would be tested by the second recall probe (Figure 1). For the remainder of this report we will refer to trials when the *relevance cue* designated the “<First>” or “<Second>” sample stimulus as “*Bound*” trials (because they entailed prioritizing a feature that, when perceived in the sample display, was bound to another feature as part of a 2-D compound stimulus), and trials when the *relevance cue* designated the “<Color>” or “<Direction>” of the two sample stimuli as “*Unbound*” trials.

### Experimental Procedure

#### Stimuli

All sample stimuli comprised 400 dots (0.08 in diameter) displayed within an invisible circular aperture (7.75° in diameter). Delayed-recall tasks featured 1-dimensional trials: on direction trials, gray dots (*L*=38, *a*=0, *b*=0) moved with 100% coherence at a constant speed of 3.2°/s in a direction that was randomly sampled (without replacement) from a list of 180 vectors that spanned the full space of 360° in increments of 2°. The direction recall interface was a dial: a white circle (7.75° in diameter) presented centrally with a white radius line (.05° wide) extending from the center to the edge of the circle (like the “needle” of an analog speedometer). On each trial, the initial angle of the needle was determined randomly, and it could be made to rotate in a clockwise or counterclockwise direction by movement of a trackball.

On color trials, the dots were stationary, and appeared in a color that was randomly sampled (without replacement) from of a list of 180 that spanned the full color space of 360° in increments of 2°. This color list was generated from an evenly distributed circle on the CIE L*a*b color space, centred at *L* = 80 with radius 60. All colors had an equal luminance and brightness and only varied in hue. The color recall interface was a horizontal bar (12.14° × 1.55°) appearing at the centre of the screen, its color transitioning smoothly across all possible colors in the color space, and a superimposed vertical white line (0.78° long, .05° wide; like the analog “tuning bar” on a radio). On each trial, the initial position of the tuning bar was determined randomly, and it could be made to translate horizontally along the color bar, to the left or to the right, by movement of the trackball.

### Behavioral tasks

Scanning was performed across three sessions, each on a different day: Day 1 -- 16 15-trial scans/blocks of 1-item delayed recall; Day 2 – 8 15-trial scans/blocks of 1-item delayed recall (for a total of 360 trials of delayed recall) plus 6 10-trials scans/blocks of MSR; Day 3 -- 10 10-trials scans/blocks of MSR (for a total of 160 trials of MSR). Before the start of each scanning session, subjects were given instructions, and the tasks were practiced both outside and inside the scanner. The task procedures in each phase are described in detail below.

#### Delayed recall

Each trial started with central presentation of a white fixation cross (0.78° width and height) against a black background (2 sec), followed by the central presentation of the sample (1 sec). The sample was either a patch of coherently moving gray dots or a patch of static dots all presented in a uniform color. The white cross returned during the subsequent 9-sec delay period, after which recall was prompted during a 3-sec window. On direction trials, subjects were instructed to click a response key when they had rotated the needle to an angle that matched their memory of the direction of motion of the sample dots. For both trial types, the needle/tuning bar became thicker (0.13°) when the response was registered, and remained at the selected position for the remainder of the 3-sec response window, followed by feedback (1 sec; errors ≤ 15° elicited “great”, > 15° and < 30° elicited “good”, and ≥ 30° elicited “poor”), followed by a gray fixation cross displayed throughout the 8-sec intertrial interval (ITI). Trial type was randomly interleaved across the whole experiment.

#### Multiple Serial Retrocuing

In the MSR task, sample stimuli were 2-D compound objects that combined the features of the delayed recall stimuli: patches of moving, colored dots. After two samples were presented, a *relevance cue* indicated what would be the critical to-be-remembered information for that trial: cues indicating “<First>” or “<Second>” designated one of the two initially presented stimuli (i.e., the “*Bound*” condition); whereas cues indicating “<Color>” or “<Direction>” designated one of the two initially presented features (i.e., the “*Unbound*” condition). Then, the remainder of all trials unfolded with two serially occurring sequences of *prioritization cue*-delay-probe (Figure 1), with each prioritization cue indicating which feature would be tested by the ensuing recall probe. The logic was that memory load was equated across both conditions - four feature tokens were initially presented, then the *relevance cue* indicated which two of these four were relevant for the trial, presumably reducing the memory load to two feature tokens - and the factor of principal theoretical interest was whether the two trial-relevant feature tokens were bound together in the same object or were drawn from two discrete objects. (Note that in this design, the factor boundedness is confounded with category homogeneity, in that *Bound* trials always required memory for a color and a direction, whereas *Unbound* trials always required memory for two colors or for two directions. However, because previous studies have shown a drop-to-baseline of the MVPA decodability of the UMI regardless of whether the two (unbound) memory items are drawn from the same or from different categories (LaRocque et al., 2013; Larocque et al., 2017; Lewis-Peacock & Postle, 2012), this confound was deemed unlikely to complicate the interpretation of the results.)

Whereas *relevance cues* presented a single word displayed in brackets -- “<First>” or “<Second>” for *Bound* trials; “<Direction>” or “<Color>” for *Unbound* trials -- *priority cues* used the same four words but without brackets. Note that the same word could never appear as both types of cue on the same trial (i.e., after a *relevance cue* of “<First>” or “<Second>”, the subsequent *priority cues* could only be “Direction” or “Color”, and vice versa). In both conditions, *priority cue 2* was equally likely to cue the feature token that had or that had not been cued by *priority cue 1*, resulting in 20 “stay” trials -- in which the same feature token was probed twice -- and 20 “switch” trials, per cell in our design. The total duration of a trial was 52 seconds, with subjects performing 160 trials in randomized order across 16 blocks of 10 trials each. All stimulus parameters were the same as in the delayed-recall task unless specified otherwise, and trial timing is illustrated in Figure 1. Both feature dimensions were randomly drawn (with replacement) from the full 360° of their respective feature spaces, in increments of 1°. The feature orientations of the second sample were constrained to a minimum angular separation of 40° relative to the first sample.

### Behavioral Data Analysis

Performance was assessed using a descriptive approach and a model-fitting approach. For both, a continuous measure of error for each response was obtained as the angular distance between the reported feature orientation and the true feature orientation. For the descriptive approach, a precision measure was then calculated as the reciprocal of the standard deviation of the error (calculated with Fischer’s formula with a correction for systematic underestimation as outlined in Bays, Catalao, & Husain (2009); http://paulbays.com/). The descriptive precision measures for each of the two probes were then submitted to a 2 × 2 repeated-measures analysis of variance (ANOVA) with category selected by the *relevance cue* (*Bound* or *Unbound*) and feature dimension (direction or color) as within-subjects factors. Trials on which no responses were given were excluded from the analyses (3%). An alpha level of .05 was applied and Bonferroni correction was used on multiple tests to control for false-positives in post-hoc testing.

The model-fitting analysis used a mixture model that decomposes the sources of error into a mixture of Gaussian variability around the target color (*P*_*T*_), a Gaussian variability around the non-target (*P*_*NT*_; also referred to as “misbinding” or “swap” errors), and a fixed probability of random guessing (*P*_*U*_). This model applied to the current data set can be described as follow:

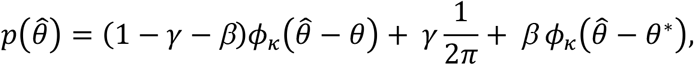

where *θ* is the target feature (probed), 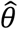 the value reported by the subjects, and *ϕ*_*κ*_ the von Mises distribution (circular analog of the Gaussian) with a mean of zero and concentration parameter *κ*. The probability of misremembering the target feature is *β* with the orientation value of the non-target feature. The probability of random guessing is captured by *γ*. Maximum likelihood estimates of the parameters *κ*, *β*, *γ* were separately obtained for each subject in each category and feature dimension conditions only for the first probe. The model components were then independently subjected to a univariate repeated-measures ANOVA with *relevance cue* (*Bound* vs. *Unbound*) and feature dimension (color vs. direction) as a within-subjects factor. Mixture modelling was only applied to the data from *probe 1* because there was not a sufficient number of trials per condition to fit the model to the *probe 2* data (due to the additional factor of “stay” vs. “switch”).

### Data Acquisition and Preprocessing

Whole brain images were acquired with the 3 T MRI scanner (Discovery MR750; GE Healthcare) at the Lane Neuroimaging Laboratory at the University of Wisconsin-Madison. High-resolution T1-weighted images were acquired for all subjects with an FSPGR sequence (8.132 ms time repetition (TR), 3.18 ms time echo (TE), 12° flip angle, 156 axial slices, 256 × 256 in-plane, 1.0 mm isotropic). Blood oxygen level-dependent (BOLD)-sensitive data were acquired using a gradient-echo, echoplanar sequence (2 s TR, 25 ms TE) within a 64 × 64 matrix (39 sagittal slices, 3.5mm isotropic).

#### fMRI data analysis

fMRI data analysis was performed using the Analysis of Functional NeuroImages (AFNI) software package (http://afni.nimh.nih.gov; Cox, 1996). All volumes were spatially realigned to the final volume of the final functional run using rigid-body realignment. The processing pipeline included slice time correction, detrending, conversion to percent signal change.

#### Generation of ROIs

Regions of interest (ROIs) were generated as a conjunction of anatomically and functionally defined voxels.

First, anatomical ROIs were generated using the Talraich anatomical atlas (TTatlas; https://sscc.nimh.nih.gov/afni/doc/misc/afni_ttatlas/index_html0). Coordinates for relevant gyri in the TTatlas were used to generate masks for each gyrus, which were then warped into an individual’s native space, and aggregated to create three regional masks. The frontal anatomical mask comprised the precentral, anterior cingulate, inferior frontal, middle frontal, superior frontal, and medial frontal gyri. The parietal anatomical mask was similarly generated and comprised the posterior cingulate gyrus, precuneus, inferior parietal and superior parietal lobules. Importantly, this included the intraparietal sulcus. The ventral occipitotemporal (VOT) mask comprised the lingual, fusiform, inferior occipital, inferior temporal, middle occipital, superior occipital gyri, and the cuneus.

Next, we fit a general linear model (GLM), separately for each subject, to the data from the delayed-recall task. Regressors of interest were delta functions placed at the beginning of stimulus onset, and a nine second boxcar modeling the delay period, all convolved with a canonical hemodynamic response function. Nuisance covariates modelled head motion and block effects. From the solution of the GLM we extracted, from each anatomical region, the top 400 voxels with the highest positive t-statistic associated with each of several contrasts: [Sample_color_ – baseline], [Sample_direction_ – baseline], [Delay_color_ – baseline], and [Delay_direction_ – baseline] to construct “feature-selective” ROIs; [(Sample_color + direction_) – baseline] and [(Delay_color + direction_) – baseline] to construct “feature-nonselective” ROIs. Of the resultant functionally defined ROIs, different instantiations would be most suitable for different analyses.

### Multivariate Inverted Encoding Modeling

Conceptually, an IEM effects a projection of the data from a large number of individual sensors into a single population-level, distributed representation (Brouwer & Heeger, 2009). To implement it, we followed the steps laid out in Ester, Sprague and Serences (2015), first, for each feature dimension (color and direction), modeling the response of each voxel as a linear sum of nine hypothetical information channels which, taken together, spanned the full stimulus space for each feature. This relationship can be expressed in the form of the following general linear equation:

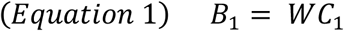

where *B*_*1*_ (*v* voxels × *n* trials) is the observed BOLD response in each voxel in each trial, *C*_*1*_ (*k* channels × *n* trials) reflects the expected responses for each information channel on each trial. For each feature, a basis set of nine modified von Mises functions (equation 4 below) was generated, each centered (by varying the *μ* parameter) around one of 9 orientations, each 40 degrees apart (at 20°, 60°, 100° through 340°), so as to cover the full 360-degree feature space. Each basis function can be construed a channel in stimulus-representation space. We set the *α*, *k*, and *β* parameters to 1,7, and 0 respectively to best approximate tuning properties of MT neurons (Duijnhouwer, Noest, Lankheet, Berg, & Wezel, 2013). To model color, we extracted a circular portion of *L*A*B* color space using the procedure similar to that outlined in Brouwer and Heeger (2009).

The first step in implementing the model (the training phase) is to regress a portion of the voxel data *B*_1_ (*v* voxels × *m* trials) against the basis set, using ordinary least-squared regression (Equation 2), to generate a weight matrix:

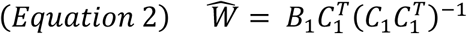

This weight matrix *W* (*v* voxels * *k* channels) constitutes a mapping from “channel space” to “voxel space”. Thus, *W* represents the relative strength (or weight) of the contribution each channel makes to the voxel’s overall response. This set of weights is sometimes referred to as a population receptive field or a voxel tuning function.

In the second step (the testing phase), the weight matrix is inverted, such that it now constitutes a mapping from “voxel space” to “channel space”, and applied to the remaining voxel data, *B*_2_ (*v* voxels * *m* trials), in order to generate an estimated representation in channel space, *C*_*2*_ (*k* channels * *m* trials):

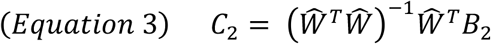

As an initial validation step, we implemented a leave-one-run out cross-validation procedure where, for each fold, one of the runs from the delayed-recall task was set aside and each time point from the remaining runs was used to generate a weight matrix for each feature dimension (color and direction) within each ROI. We then inverted the weight matrix and applied it to data from the left-out run to generate reconstructions in channel space (also referred to as “channel tuning functions” CTFs). Reconstructions from each iteration of the leave-one-run-out procedure were then aligned and averaged together to generate reconstructions for the delayed-recall task, which we then quantified using the procedure outlined below. These results are shown in Figures 4, 5 and 6.

**Figure 3.**
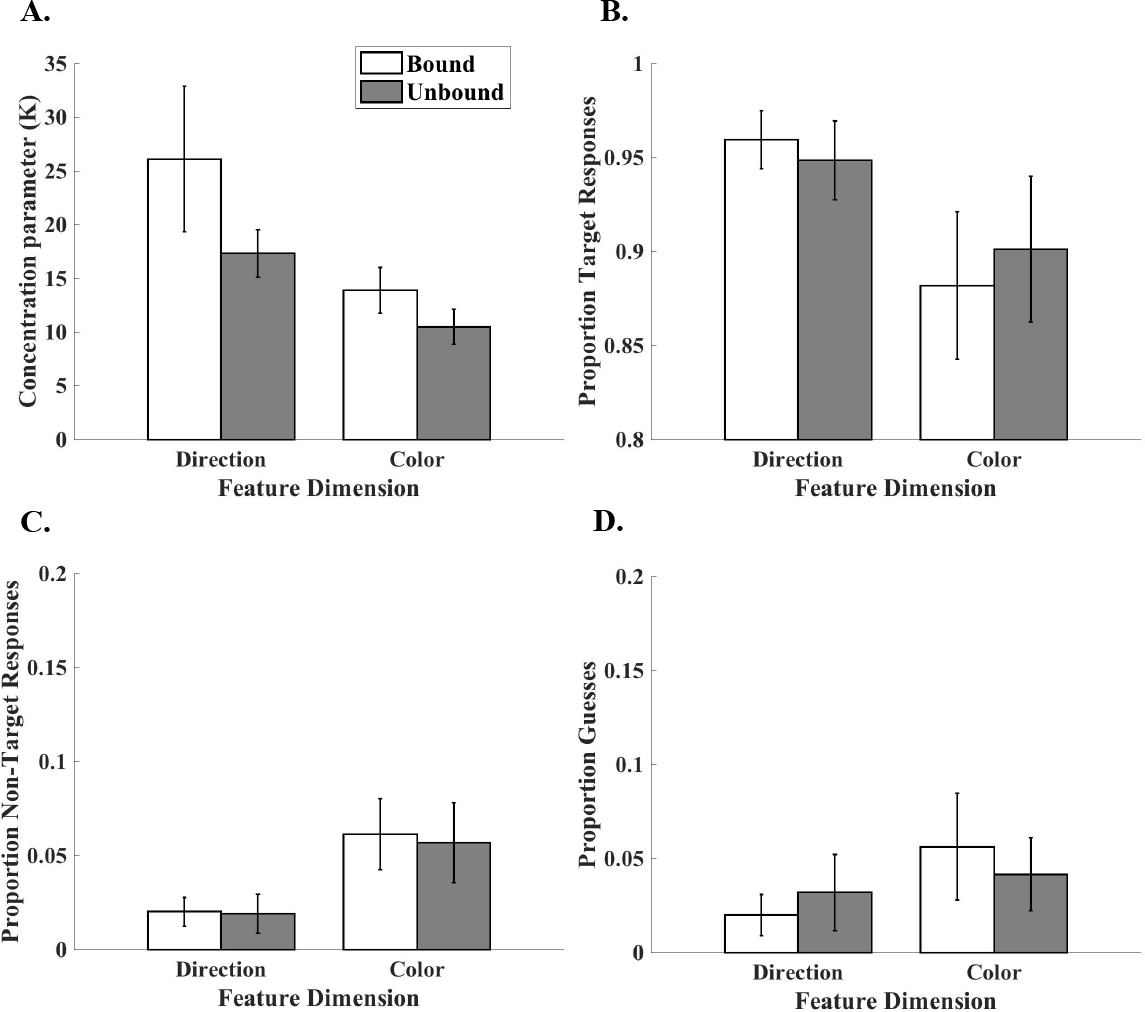
Mixture-model estimates for *probe 1* responses: (A) Concentration parameter; (B) probability of response to target; (C) swap errors; and (D) guesses.

**Figure 4.**
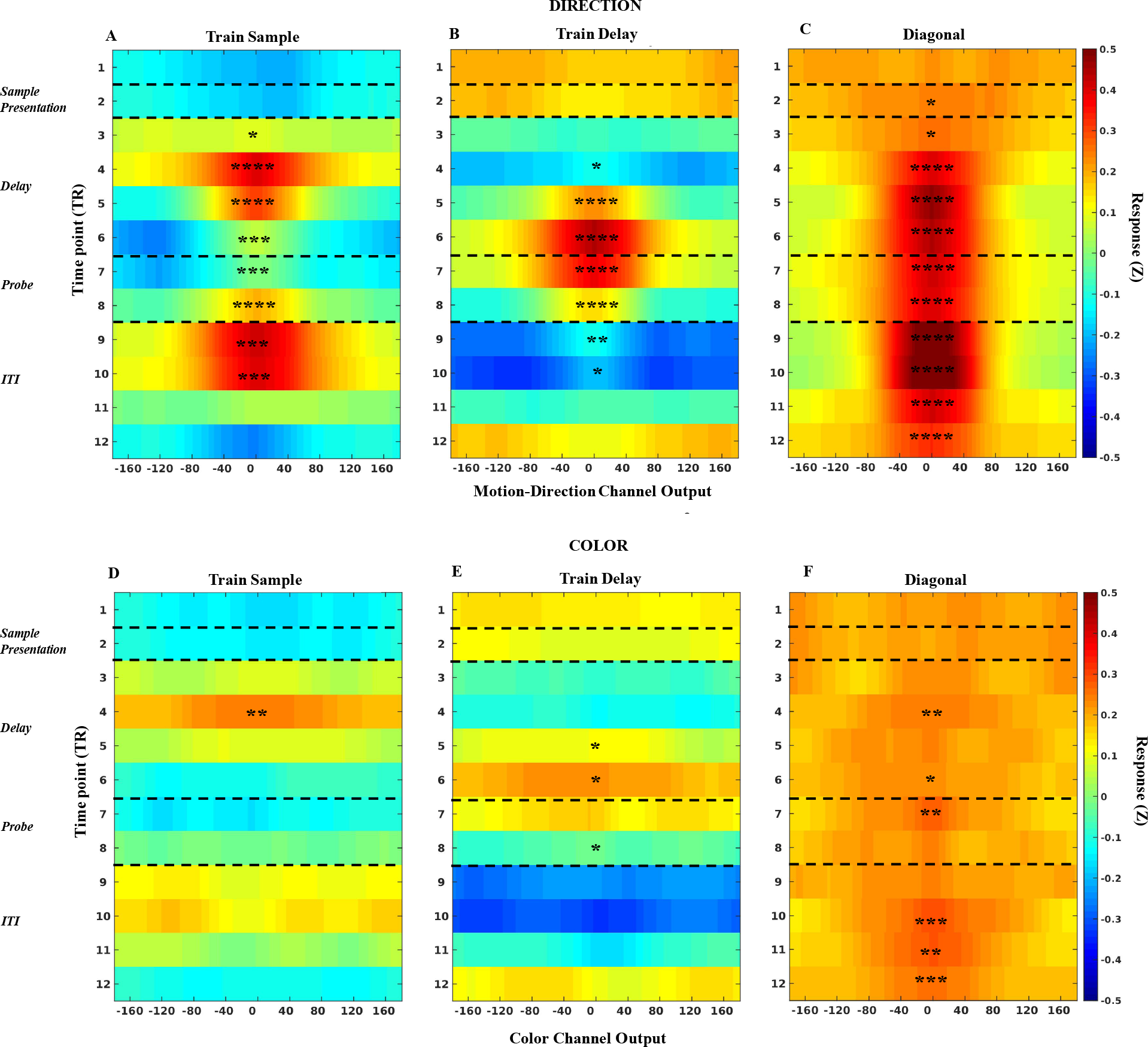
Time courses of IEM reconstructions from the 1-item delayed-recall task, in feature-selective VOT ROIs. Rows illustrate results for direction (top, panels (A), (B), and (C)) and color (bottom, panels (D), (E), and (F)), and columns the results for three procedures: train on TR 4 (a “perception/encoding” model) and test at every TR (left, panels (A) and (D)); train on TR 6 (a “delay” model) and test at every TR (middle, panels (B) and (E)); and train and test at each TR (“along the diagonal,” right, panels (C) and (F)). In each panel, channel is arrayed along the horizontal axis, from −160° to 160°, time (in TRs) proceeds from top to bottom, and the dependent data are channel responses (averaged after aligning each trial to 0°). Significance of the reconstruction at each TR, determined by bootstrapping, is indicated by asterisks (* = p<.05; ** = p<.01; *** = p<.001; **** = p<.0001).

**Figure 5.**
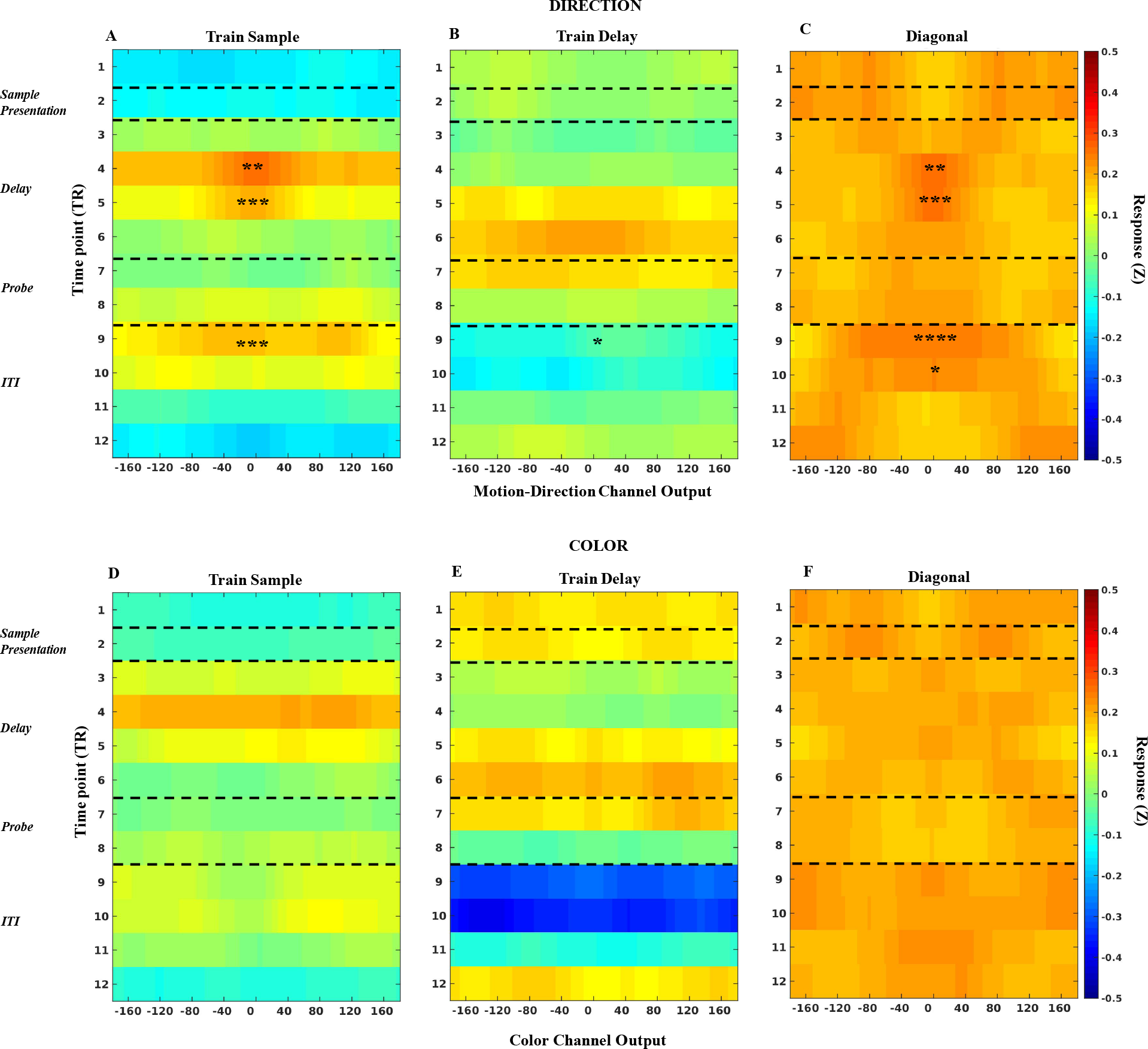
Time courses of IEM reconstructions from the 1-item delayed-recall task, in feature-selective parietal ROI. All display conve

**Figure 6.**
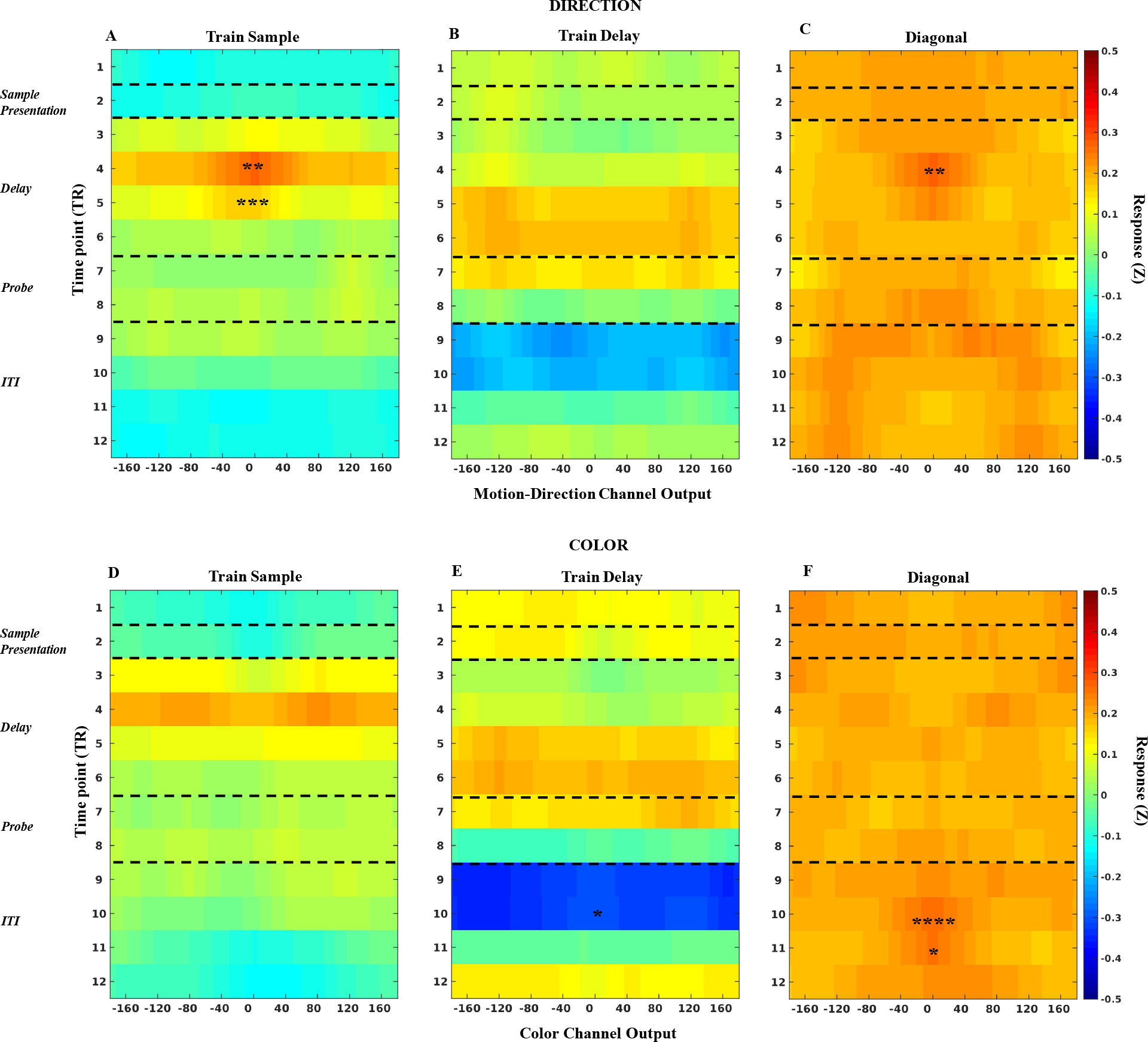
Time courses of IEM reconstructions from the 1-item delayed-recall task, in in feature-selective frontal ROIs. All display conventions the same as Figure 4.

Once the weight matrix was generated from delayed-recall data, data from each time point in the MSR task (testing phase) was multiplied by the inverted weight matrix as described in Equation 3 to generate a reconstruction time course of direction. Each of these feature-specific reconstruction time courses were then circularly shifted to a common center (0°) and averaged with those from like trials. Thus, for example, to generate the “Attended” reconstructions for the *Unbound* condition (Figure 7), channel outputs from trials for which “<Direction>” was *relevance*-cued, and for the item that was cued by *priority cue 1*, were aligned along the *priority*-cued item’s direction and averaged together. In order to generate the smooth, 360-point functions shown in Figures 8 and 10, we repeated the IEM analysis a total of 39 times and shifted the centers of the direction or color channels by 1° on each iteration.

**Figure 7.**
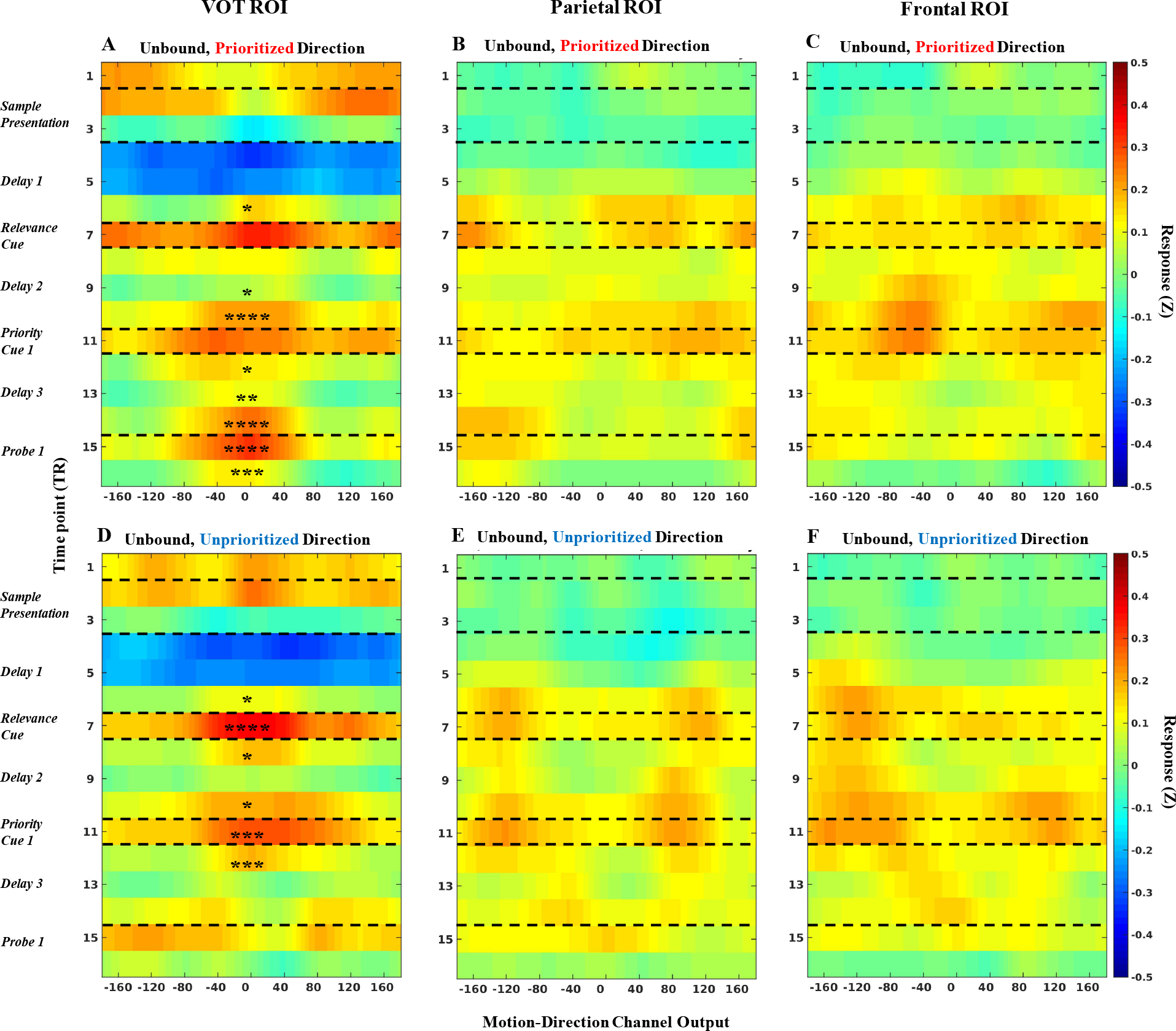
Time course of the neural representation of direction during *Unbound* trials, across the first half of the MSR task, as a function of priority status. The top row of panels illustrates the reconstruction of the direction of motion was cued by *priority cue 1*, and the bottom row illustrates the reconstruction of the direction of motion was not cued by *priority cue 1*. All display conventions within each panel are the same as Figure 4.

**Figure 8.**
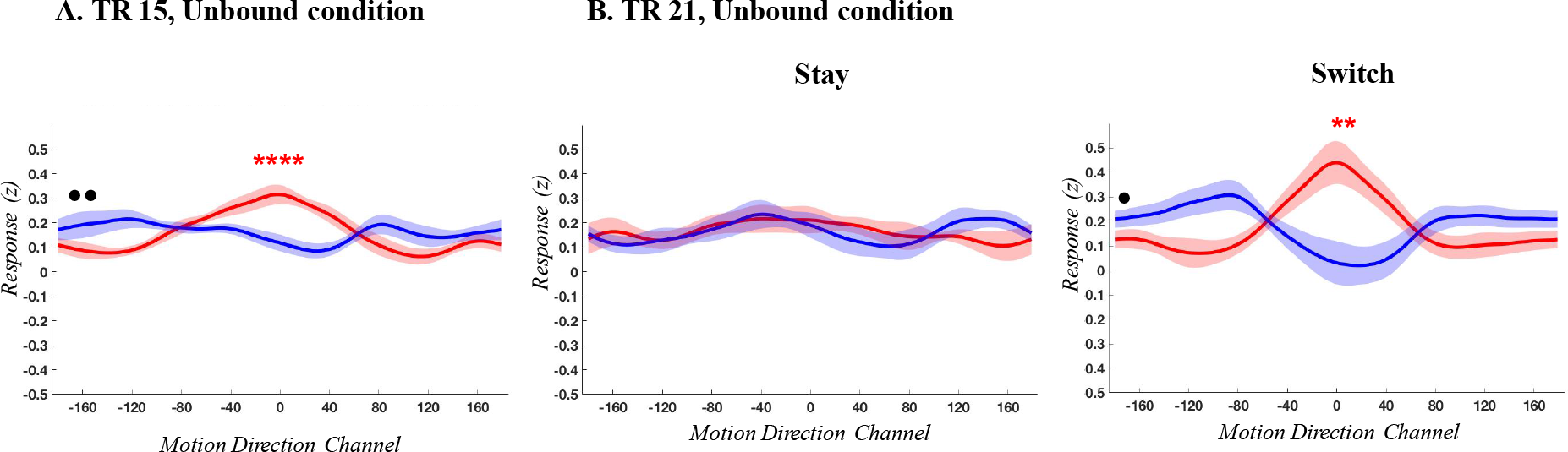
The reconstruction of the neural representation of direction of the PMI (red) and the UMI (blue) from individual TRs of *Unbound* trials, in the feature-selective VOT ROI, at (A) TR 15 (immediately prior to *probe 1*), (B) TR 21 (immediately prior to *probe 2*) on “stay” trials, and (C) TR 21 on “switch” trials. In each plot, channel is arrayed along the horizontal axis, and the dependent data are channel responses (averaged after aligning each trial to 0°). The width of each reconstruction trace represents the standard error of the mean across subjects, interpolated across the 360 discrete data points in the averaged data. Significance of the reconstructions of the PMI and UMI is indicated by red and blue asterisks, respectively (* = p<.05: ** = p<.01; *** = p<.001; **** = p<.0001). Significance of the difference between the baseline parameter of the of the PMI vs. the UMI reconstruction are indicated with black circles (• = p<.05; •• = p<.01).

**Figure 9.**
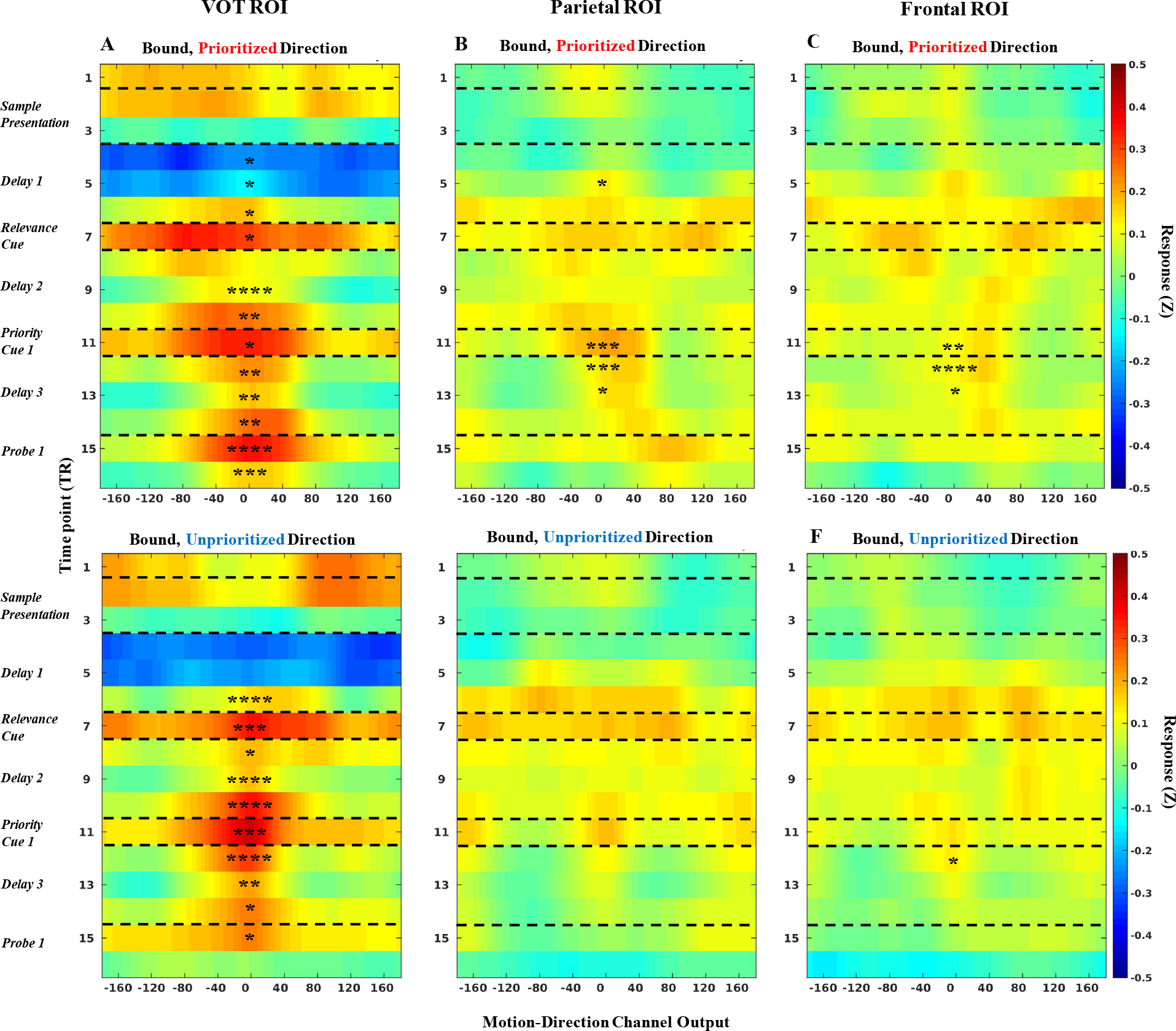
Time course of the neural representation of direction during *Bound* trials, across the first half of the MSR task, as a function of priority status. All display conventions within each panel are the same as Figure 4. The top row of panels illustrates trials during which direction became the PMI during *delay 3*, and the bottom row illustrates trials during which direction became the UMI during *delay 3*. All display conventions within each panel are the same as Figure 4.

**Figure 10.**
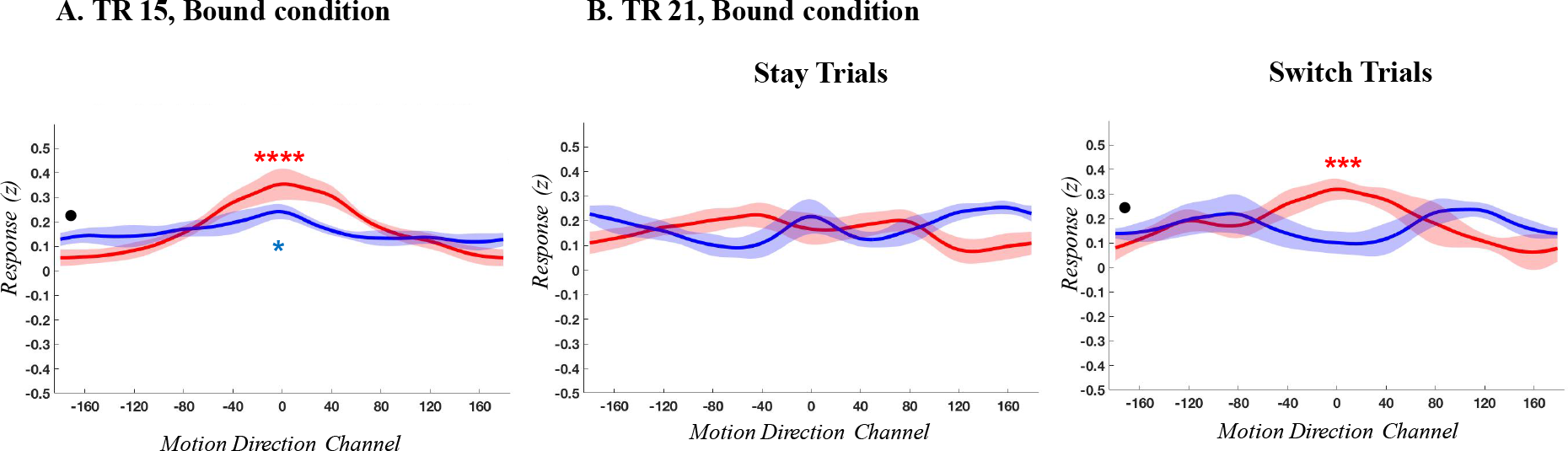
The reconstruction of the neural representation of direction of the PMI (red) and the UMI (blue) from individual TRs of *Bound* trials, in the feature-selective VOT ROI, at (A) TR 15 (immediately prior to *probe 1*), (B) TR 21 (immediately prior to *probe 2*) on “stay” trials, and (C) TR 21 on “stay” trials. All display conventions the same as Figure 8.

Reconstructions were then quantified using a bootstrapping procedure similar to Ester et al. (2015). In each ROI, for each feature dimension, each time point, and each condition, reconstructions from all nine subjects were randomly sampled with replacement nine times to generate a 9×360 dimension resampled reconstruction matrix. This resampled matrix was averaged across the first dimension (subjects), yielding an averaged reconstruction that was then fit with the following von Mises response function:

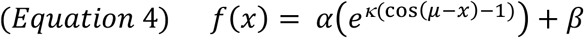

where *x* is a vector of 360 channel responses. *μ*, *κ*, and *β* correspond to the center (i.e., mean), concentration (i.e., inverse of width) and baseline (i.e., vertical offset) of the function, respectively, and *α* corresponds to the amplitude of the function (i.e., vertical stretching/scaling). Fitting was performed using a combination of a gridsearch procedure and ordinary least squares regression. The response function (Equation 4) was defined by setting the *μ* parameter to 0 (reflecting our alignment across trials to a 0° centered channel) and the *β* parameter to 1. A range of plausible *κ* values was then defined (from 1-30 in increments of 0.1). For each *κ* value, a design matrix was then generated containing the response function and a constant term (i.e., a vector of 1s). Fitting the reconstructed feature tuning curves yielded estimates of *α* and *β*, the regression coefficients for the design matrix and constant terms respectively. The best fitting curve that minimized the sum of squared errors between response function and data was then selected.

This procedure was repeated 10,000 times in total, yielding 10,000 bootstrapped estimates of amplitude, baseline, and concentration. To test the whether the amplitudes of the PMI and UMI representations were significantly above baseline levels, (one-tailed) *p*-values for the robustness of the PMI and UMI feature reconstructions were separately calculated in the *Unbound* and *Bound* conditions by assessing the percentage of bootstrapped iterations whose amplitude estimates were negative. In other words, statistical significance at an alpha level of 0.05 implies that at least 95% of resampled reconstructions have a positive amplitude (*p*_pos_).

In order to test whether feature reconstructions of PMIs were stronger than those of UMIs, we computed the difference between the bootstrapped amplitudes of the PMI and UMI reconstructions, forming a distribution of difference scores. We assessed the percentage of bootstrapped iterations whose amplitudes were negative. In other words, statistically significant differences (at *p*<0.05) between the PMI and UMI conditions would indicate that 95% of the differences in the resampled amplitude estimates of the PMI and UMI reconstructions were positive (*p*_pos_). The difference between the PMI and UMI reconstructions were separately calculated in the *Unbound* and *Bound* conditions. The same principle of statistical testing was applied to the baseline parameter. We were particularly interested in the attentional modulations of feature representations late in the delay period, namely in the time points 15 and 21 in response to *priority cue 1* and *priority cue 2*, respectively. Therefore, the tests in the results sections are mainly focused on these time points (see figures 8 and 10). However, reconstructions of the entire time courses of the delay periods of both priority cues are presented in supplementary figures along with the statistics on the amplitude and baseline differences between PMI and UMI feature reconstructions.

*Delay 4*, following the presentation of *priority cue 2*, differed from delay 3 because only in the former could the uncued feature be dropped from WM. The results from *delay 4*, therefore, could provide insight about whether the putative “dropping” operation differs as a function of boundedness^1^. Because analyses of data from *delay 4* could only use half of each subject’s data, due to only switch trials affording a comparison of the transition of an item from PMI to UMI/”dropped memory item” (DMI) following *priority cue 1* vs. *priority cue 2*, we restricted our analyses of *delay 4* of MSR trials to TR 21, the final TR expected to reflect delay-period processing uncontaminated by the processing of *probe 2* (see, e.g., the BOLD time courses in Figure 1), and carried out the same analyses on TR 15, to allow for like-to-like comparison of stimulus feature representations between *delay 3* and *delay 4*. Over the course of data analysis, we observed that, in some conditions, the reconstruction of the UMI produced amplitudes that were negative (c.f., (van Loon et al., 2018)). Because these outcomes were not predicted, we evaluated whether these “negative reconstructions” differed significantly from 0 by calculating two-tailed *p*-values: first, the probabilities of obtaining a positive (*p*_pos_) or a negative (*p*_neg_) amplitude among the 10000 bootstrapped amplitudes were calculated; next, two-tailed *p*-values were obtained using:

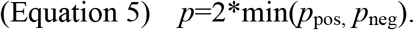

### Multivariate pattern analyses

We carried out multivariate pattern analyses (MVPA) to clarify and/or refine the interpretation of some of the findings from the IEM analyses, using L2-regularized logistic regression (with a lambda penalty term of 25) applied to z-scored BOLD, and implemented with the Princeton Multi-Voxel Pattern Analysis toolbox (www.pni.princeton.edu/mvpa/) and custom scripts in MATLAB (c.f., Lewis-Peacock et al., 2012). MVPA was carried out on two levels of stimulus information: (i) within feature (color and direction labeled as belonging to one of four quadrants in their respective 360° stimulus spaces, carried out in feature-selective ROIs); and (ii) between feature (color vs. direction), carried out in feature-nonselective ROIs).

To assess the representation of stimulus-level of information, we trained two classifiers, one from color-sensitive voxels and one from motion direction-sensitive voxels as in the IEM analyses, separately for each ROI, to classify motion-direction and color values categorized as belonging to one of four quadrants (centered at 45°, 135°, 225° and 315°, each spanning a 90° wedge of positions within the full 360° range of possible colors and motion directions). Categorizing stimuli in this way enabled us to determine whether coarse stimulus information might be decodable in the frontal and parietal ROIs for which the IEMs failed to reconstruct exemplar-specific feature information. Classifiers were trained on late delay-period data from the delayed-recollection task (TR 6), with k-fold cross-validation (train on 23 runs, test on the 24^th^), classification accuracy averaged across folds and compared against chance performance with two-tailed *t*-tests (Bonferroni corrected).

Additionally, to assess the representation of higher-order information about stimulus category, we trained classifiers (in feature-nonselective ROIs) to accurately distinguish between the categories of color vs. direction on data from the 1-item delayed recall task. MVPA methods were the same as described above, with the exception that on two labels were used for training (“color,” “direction”), and so to statistical significance was assessed with two-tailed *t*-tests comparing accuracy to chance performance (50%).

Finally, to assess evidence for cognitive control-related activity, we also applied the category-level decoders trained on data from the 1-item delayed-recall task to data from the MSR task. Because a hallmark of control is that it should dynamically track changing contingencies within individual trials, we carried out these analyses by applying late-delay classifiers from the 1-item delayed-recall task to every TR of “switch” trials from the *Bound* condition of the MSR task. This would generate classification time courses for MSR trials that featured within-trial switches of priority between stimulus category. For this analysis, at each time point of the MSR task, and for each category, a measure of pattern similarity was computed between the voxel patterns for that TR and the late-delay patterns from the 1-item delayed-recall task. Using logistic regression, each category’s pattern similarity score was then converted into “classifier evidence,” a value between 0 and 1 that can be interpreted as the extent to which the pattern at the tested TR matches the pattern learned by the classifier (i.e., conceptually similar to a correlation coefficient; c.f., Lewis-Peacock & Postle, 2012; Polyn, Natu, Cohen, & Norman, 2005). Average classifier estimates were computed by sorting trials according to the feature dimension selected by *priority cue 1* and the feature dimension selected by *priority cue 2*. Statistical significance of the evidence between the feature dimensions as function of the priority cues was computed by pairwise *t*-tests at each time-point (Bonferroni corrected).

## Results

### Behavioral

#### Model-free measures

Analysis of the precision of responses revealed only main effects of feature dimension: *probe 1* [*F*(1,8) = 10.76, *p*<.05, *η*_*p*_^*2*^=.57] with an overall higher precision for direction (*M* = 2.99, *SE* = .61) than for color (*M* = 1.44, *SE* = .19) responses (other effects n.s.); *probe 2* [*F*(1,8) = 7.62, *p*<.05, *η*_*p*_^*2*^=.49] with an overall higher precision for direction (*M* = 2.46, *SE* = .49) than for color (*M* = 1.53, *SE* = .25) responses (other effects n.s.; Figure 2).

#### Mixture modeling

Although inspection of results for *κ* (Figure 3.A.) suggests qualitatively similar patterns to those for precision, ANOVAs indicated, instead, a greater sensitivity to boundedness (main effect of *relevance cue* [*F*(1,8) = 5.66, *p*<.05, *η*_*p*_^*2*^=.41], with *κ* higher in the *Bound* (*M* = 20.02, *SE* = 2.91) than the *Unbound* (*M* = 13.94, *SE* = 1.4) condition), and the difference between feature dimensions no longer meeting the threshold for significance [*F*(1,8) = 4.84, *p*=.059, *η*_*p*_^*2*^=.38] (direction trials (*M* = 21.75, *SE* = 3.78); color trials (*M* = 12.22, *SE* = 1.50)). The interaction between boundedness and feature dimension for *κ* did not reach significance (*F*<1).

The greater difficulty of color than direction performance, as suggested by the descriptive statistics, was captured in the model’s estimates of *P*_*T*_ ([*F*(1,8) = 7.51, *p*<.05, *η*_*p*_^*2*^=.48], with a higher *P*_*T*_ for direction (*M* = .95, *SE* =.016) than for color (*M* = .89, *SE* = .036) responses (other *F*s < 1)), and of *PN*_*T*_ ([*F*(1,8) = 8.17, *p*<.05, *η*_*p*_^*2*^=.51], with a lower *PN*_*T*_ for direction (*M* = .02, *SE* =.006) than for color (*M* = .059, *SE* =.016) (other *F*s < 1)). There were no differences in *P*_*U*_ (*F*_s_ < 4).

### IEM Reconstructions

#### 1-item delayed recall

##### Sample-evoked ventral occipitotemoral ROI

In the sample-evoked VOT ROI, stimulus feature reconstructions were markedly superior for direction than for color. For direction, a model that was trained on TR 4 (i.e., the volume collected from 5-6 sec after sample onset, when the sample-evoked BOLD response was expected to be maximal (c.f., Figure 1.C.)) yielded a robust reconstruction when tested at that same TR (with leave-one-run-out k-fold cross validation). Furthermore, sweeping this model across all 12 TRs of the trial yielded reliable reconstructions spanning from the first TR of the delay period through to the second TR after the response window, thereby demonstrating robust cross-temporal generalization, and indicating that stimulus direction was represented, in part, with a perceptual neural code throughout the trial (Figure 4.A.). The same qualitative pattern of reconstruction was observed when this process was repeated with a model trained on data from TR 6, which was intended to capture signal primarily attributable to delay-period processing (Figure 4.B.) Finally, direction reconstruction was achieved at all but the TR preceding sample presentation when a model was trained at tested at each TR (i.e., “along the diagonal” of a cross-temporal matrix, Figure 4.C.). For color, an IEM could be successfully trained at TR 4, but did not generalize to any other TRs (Figure 4.D.), and one trained at TR 6 generalized to TRs 5 and 8 (Figure 4.E.). When models were trained and tested at each TR, the reconstruction of color information was successful for only a subset of TRs associated with the perception/encoding and retention of color information, as well as for TRs during the ITI that followed the probe (Figure 4.F). Although one might expect poorer IEM reconstruction for the stimulus feature that was remembered less well (i.e., color recall was inferior to direction recall), it could also be the case that the reconstruction of neural representations of color would have been more robust had we attempted to optimize for IEM the plane of the slice through CIE space that we selected to generate our stimuli. Additionally, our method does not allow us to know the extent to which verbalization may have contributed to color WM performance, a possibility that would not be expected to produce robust IEM in VOT ROI.

##### Delay-evoked Parietal and Frontal ROIs

In parietal cortex, successful reconstruction of sample direction was restricted to just a few TRs associated with encoding/early delay and with recall/response, and no successful reconstructions of sample color (Figure 5). In frontal cortex the pattern was similar, with the exception that there were a few successful reconstructions of sample color in ITI TRs (Figure 6).

#### Multiple Serial Retrocuing

Results from the analyses of the 1-item delayed-recall data indicated that the IEM reconstruction of the neural representation of direction was markedly superior than that for color, and, furthermore, that the reconstruction of direction was markedly stronger in the VOT than in the parietal and frontal ROIs. Therefore, to maximize the sensitivity for addressing our question of principal interest, we focused on the representation of direction in the VOT ROI by training a “1-item delay” IEM from TR 6 of the delayed-recall task and testing it on data from the MSR task.

##### Unbound condition

On trials of all types, the neural representation of direction in the VOT was robust during the delay period prior to the *relevance cue* (TR 6 in panels A and D of Figures 7 and 9). Furthermore, on *Unbound* trials, after the *relevance cue* indicated “<Direction>”, the reconstructions of the direction of the sample that would become the PMI (Figure 7.A.) and of the sample that would become the UMI (Figure 7.D.) were both robust during the TRs leading up to and immediately following *priority cue 1* (TR 11), with statistically comparable reconstructions from the data collapsed across TRs 9, 10, and 11 (two-tailed, *p*=.69).

Once *priority cue 1* designated one item the PMI and one the UMI, the amplitude of the reconstructions diverged markedly. IEM of the PMI produced robust delay-period reconstructions from TR 12 through TR 15, with the reconstruction strengthening across the delay preceding *probe 1* (*slope* = .039, *p*<.05). For IEM of the UMI, in contrast, the amplitude of the 0° channel declined significantly from TR 12 to TR 15 (*slope* =−.069, *p*<.001), and no reconstructions were reliable from TRs 13-16. Indeed, the output of the channels near 0° channel dropped below that of flanking channels, although this trend toward a significantly “negative” channel tuning function did not achieve significance when the data were collapsed across TRs 13, 14, 15 (two-tailed, *p*=0.27). Finally, statistical comparisons confirmed that reconstructions of the PMI were higher in amplitude than those of the UMI (signal from both collapsed across TRs 13 through 15, one-tailed, *p* < .0001).

No reconstructions were reliable in the parietal and frontal ROIs. Unexpectedly, however, in all ROIs, the overall magnitude of IEM channel outputs increased markedly from *Delay 1* (TR 4 through 6) to *Delay 2* (TR 8 through 10), as reflected in significant increases in the values of the baseline parameter in the VOT (*p*<.001, two-tailed) and parietal (*p*<.01, two-tailed) ROIs; for the frontal ROI (*p*=.11). (Possible interpretations of changes in baseline will be considered in the section on *Multivariate classification*, below.)

*The DMI following priority cue 2*. At TR 21, on “stay” trials, neither the PMI nor the DMI could be reconstructed (Figure 8.B.). On “switch” trials, however, the IEM reconstruction of the newly cued PMI (which had been flat and nonsignificant during the previous delay; Figure 8.A.) was robust at the end of *delay 4* (TR 21), and that of the DMI trended in the opposite direction: that is, the reconstruction displayed minimal channel output along the aligned direction and maximal output for channels representing the opposite direction. Because such a “complementary” reconstruction (in reference the complementary nature of a cosine wave in relation to a sine wave) was not predicted, we assessed its significance with a two-tailed test, which indicated that it missed the threshold for significance (*p* = .06, Figure 8.C., left-hand column). At TR 21 the PMI and DMI differed in amplitude (*p* < .01) and in baseline (*p* < .05, two-tailed).

##### Interim summary of results from Unbound condition

For trials on which the *relevance cue* indicated “<Direction>”, the subsequent *priority cue 1* influenced the representation of both features held in WM: the representation of the PMI increased in strength across the subsequent *delay 3*, whereas the representation of the UMI decreased in strength to the point that it could no longer be reconstructed by the end of *delay 3*. This is the pattern of results that would be expected if the principles of biased competition apply to the selection of one from among two working-memory representations of stimuli in the same way that they do for the selection of one from among two objects in a visual scene. On trials when *priority cue 2* prompted a switch of prioritization status, the IEM reconstruction of the newly designated PMI increased over the course of *delay 4*, whereas that of the newly designated DMI decreased over the course of *delay 4*, from being significantly positive at the beginning (Figure 7.A.) to being nonsignificant and bordering on complementary to the trained model at the end. Additionally, the baseline parameter of this complementary reconstruction of the UMI during *delay 4* was higher than that of the PMI.

##### Bound condition

After the *relevance cue* indicated “<First>” or “<Second>”, the reconstructions from the VOT ROI of the direction of the sample that would become the PMI (Figure 9.A.) and of the sample that would become the UMI (Figure 9.D.) were robust during the TRs leading up to and immediately following *priority cue 1* (TR 11), with statistically comparable reconstructions from signal collapsed across TR9, TR10, and TR11 (two-tailed, *p*=.77). Unlike in the *Unbound* condition, however, the designation by *priority cue 1* of the PMI and the UMI had only a relatively minor effect on the IEM reconstructions. Although the IEMs of the PMI reconstructions were sustained across the ensuing delay period, their strength did not increase (slope = .026, *p* = 0.25). Furthermore, although the amplitude of the reconstructions of the UMI decreased across this delay period (slope = −.050, *p* < 0.05), they remained statistically significant across the delay period, and only beginning with TR 15 did the reconstruction of the UMI decline in amplitude to a point at which it was significantly lower than that of the PMI (*p* < .05).

Results from the *Bound* condition also differed markedly from the *Unbound* condition in the parietal and frontal ROIs, in that representations of the direction of the PMI became significant with the onset of *priority cue 1* and for a few TRs into the ensuing *delay 3* (Figure 9.B. and C., top row), as well as, in the frontal ROI, of the UMI for a single TR.

The overall magnitude of IEM channel outputs increased from *Delay 1* to *Delay 2*, as reflected in the values of the baseline parameters, although, as with the *Unbound* condition, this increase only reached significance in the VOT (two-tailed, *p*<.0001) and parietal (two-tailed, *p*<.05) ROIs.

*The DMI following priority cue 2*. The patterns at late-*delay 4* (TR 21) mirrored those from the *Unbound* condition: On “stay” trials neither the PMI nor the DMI could be reconstructed (Figure 10.B.); and on “switch” trials, reconstruction of the PMI was significantly positive (*p* < .0001) and that of the DMI was nonsignificant, but trending toward a complementary reconstruction (*p* = .09, two-tailed); and the two differed significantly from each other (*p* < .0001).

##### Interim summary of results from Bound condition, and comparisons between conditions

On trials when “direction” was cued by *prioritization cue 1*, the IEM reconstruction of the PMI remained robust across the ensuing delay period, but it did not increase in strength. On trials when “color” was cued by *prioritization cue 1*, although the amplitude of the neural representation of the UMI declined across the ensuing delay period, it nonetheless remained significantly elevated throughout the delay (Figure 9). Together, these results are consistent with the idea that a cardinal principle of object-based attention may apply to visual WM in a manner similar to visual perception: When one feature of an object is selected, the benefits of attention extend to all features of that object (e.g., Duncan, 1984; Egly et al., 1994). Although we cannot rule out the possibility that the UMI may have also benefited from the allocation of spatial attention to the stimulus of which it was a part, this possibility seems unlikely because, by this account, spatial attention would also be expected to boost the strength of the DMI, but this was not observed: Following *prioritization cue 2*, the representation of the PMI at TR 21 was positive, and the representation of the DMI was nonsignificant (and, indeed, trending in a direction opposite to what a spatial attention account would predict). Because this pattern mirrored what was observed in the *Unbound* condition, the implication is that the dynamics of “dropping” a no-longer-needed feature from WM may be similar across these two conditions.

To quantify comparisons across conditions, we first established that the amplitudes of the reconstructions of direction that would become the PMI during *delay 3* (i.e., at TRs 12-15) were comparable during the time immediately preceding *priority cue 1* (i.e., at TRs 9-11, *p* = .31; n.s.). Next, comparison of the change in PMI amplitudes across *delay 3* indicated that the strengthening of the PMI that was observed in the *Unbound* condition did not differ significantly from the flat slope of the amplitude of the PMI in the *Bound* condition (slope difference = .01; two-tailed, *p* = .74). Finally, at TR 15, the amplitudes of the PMI did not differ between *Bound* and *Unbound* the conditions (two-tailed, *p* = .52). Turning to the UMI, we first established that the amplitudes of the representations of direction that would become the UMI during *delay 3* were comparable during the time immediately preceding *priority cue 1* (i.e., at TRs 9-11: two-tailed, *p* = .62; n.s.). Next, comparison of the slopes of the decline in the strength of the UMI across *delay 3* indicated that the weakening of the UMI across *delay 3* was not significantly different in the two conditions (slope difference = -.019, *p* = .55). At TR 15, however, the amplitudes of the UMI differed between *Bound* and *Unbound* conditions (two-tailed, *p* < .05). For *delay 4*, on “switch” trials, in both the *Bound* and the *Unbound* conditions the reconstructions at TR 21 were significantly positive for the PMI and trending toward complementary for the UMI. In summary, although many of the patterns observed during the *Bound* vs. the *Unbound* conditions did not differ statistically from each other, the difference in the amplitude of the UMI at the end of *delay 3* confirmed the theoretically important observation that the processing of the UMI is dependent on the effects of object-based attention.

### Multivariate pattern classification

#### Delayed-recall task

In the VOT ROI, successful decoding of direction (*t(8) = 3.45, p*<.05; Figure 11.A.) validated the logic of decoding stimulus identity by binning specific items into arbitrarily defined quadrants. It also argued against the possibility that the failure to successfully reconstruct direction in parietal and frontal ROIs (*t*s < 1.36), as well as the failure to reconstruct color in any of the ROIs (*t*s < 1.03; Figure 11), might be attributable to the method, per se. Indeed, these null decoding findings were broadly in line with the results of the IEM analyses. In contrast to these null findings, successful decoding of trial type (color vs. direction) in parietal (*t*(8)=10.61, *p*<.001) and frontal (*t*(8)=8.4, *p*<.001) regions, (as well as in VOT cortex (*t*(8)=6.72, *p*<.001); Figure 11.B.) indicated that these regions tracked higher-order task-related information.

**Figure 11.**
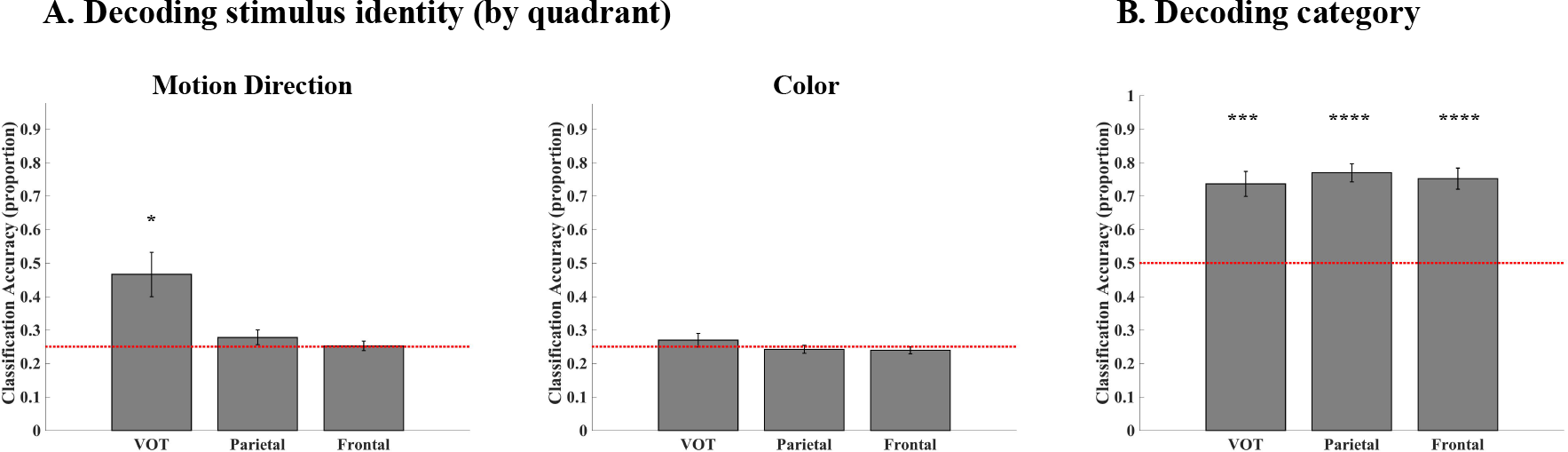
MVPA decoding of stimulus feature information from the late delay period of the 1-item delayed-recall task. (A) Performance of classifiers trained on data labeled according to post hoc-defined quadrants in 360° stimulus space. (B) Performance of classifiers trained on data labeled according to feature category (i.e., color vs. direction). Statistically reliable classification is denoted by asterisks (* = p<.05: ** = p<.01; *** = p<.001; **** = p<.0001).

#### Multiple serial retrocuing task

##### MVPA of stimulus category

Although parietal and frontal cortex are generally associated with the control of WM (e.g., Brincat, Siegel, Nicolai, & Miller, 2017; Pribram et al., 1964), for the successful decoding of trial type (e.g., Figure 11.B.) to be interpreted as reflecting control-related activity, one would want to see that it dynamically tracks changing contingencies within individual trials. We assessed this possibility by applying late-delay classifiers from the 1-item delayed-recall task to every TR of “switch” trials from the *Bound* condition of the MSR task, so as to generate classification time courses. (Note that only the *Bound* condition included within-trial switches between stimulus category.) These analyses, carried out in feature-nonselective ROIs in parietal and frontal cortex, revealed that information about color and direction were represented, to the same extent, from the beginning of the trial until TR 11 (the onset of *prioritization cue 1*). At this point in the trial, classifier evidence for the cued feature increased steeply and evidence for the uncued feature decreased steeply. At TR 17 (the onset of *prioritization cue 2*, a “switch” cue in these analyses), these patterns reversed, with evidence for the newly prioritized feature rising precipitously, and evidence for the newly unprioritized falling precipitously (Figure 12.A.). Such patterns of event-locked shifts of category representation are consistent with a role for parietal and frontal regions in the control of stimulus representation and of behavior on this task.

**Figure 12.**
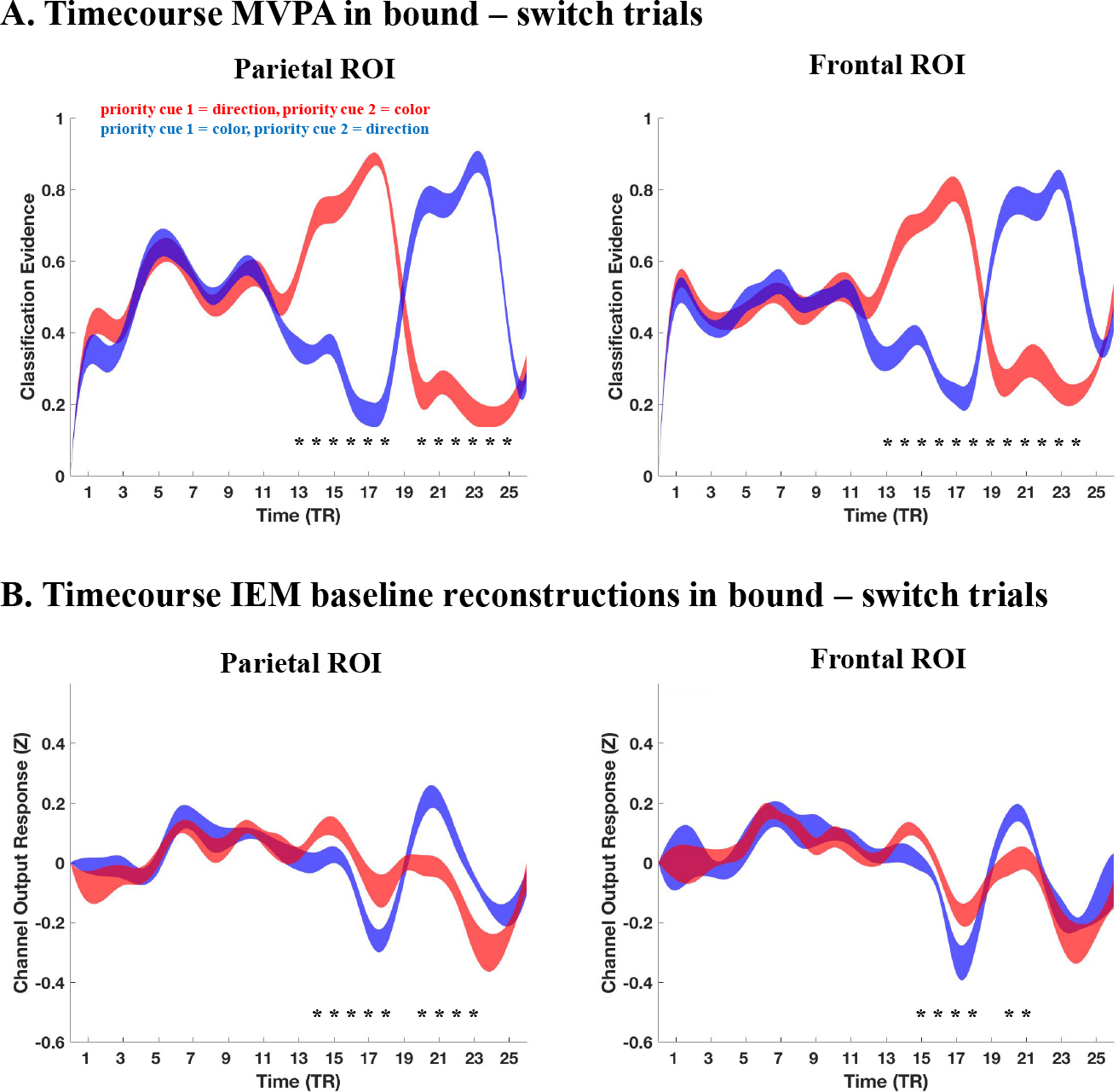
The time-course of model performance on “switch” trials from the *Bound* condition of the MSR task. (A) Trial-averaged time courses of MVPA evidence for the representation of direction of motion, on trials when motion was cued by *priority cue 1* (red) vs. trials when color was cued by *priority cue 1* (blue). (B) Trial-averaged time courses of the value of the baseline parameter from IEM of the same trials from panel A. Asterisks indicate TRs for which values from the two trial types differ significantly; the width of each trace represents the standard error of the mean across subjects.

##### Timecourse of the baseline parameter from IEM mirrors MVPA of stimulus category

Inspection of the IEM timecourses of the first half of the MSR trials (Figures 7 and 9), as well as the late-delay reconstructions from *delay 4* (Figures 8 and 10) reveals considerable variation in the baseline parameter of IEMs. By definition, this parameter does not relate directly to stimulus representation. To explore the possibility that these patterns of variation may index control-related activity, we plotted the values of the baseline parameter from tests of the late-delay IEM of 1-item delayed recall on each TR of the trials of the MSR task featured in Figure 12.A. As illustrated in Figure 12.B, the fluctuations of the baseline parameter closely followed those of the MVPA of feature category.

#### Interim summary of MVPA results

The results of MVPA of delayed-recall data binned post hoc into quadrants in stimulus-feature space were broadly consistent with the IEM results, in that evidence for stimulus-level representation of sample information was only reliably found for the dimension of direction-of-motion, and only in the VOT ROI. This reinforces the idea that stimulus-specific direction information was most prominently represented in VOT cortex. Classification at the more abstract level of stimulus category (i.e., color vs. direction), however, was reliable in all three ROIs. Furthermore, when the late-delay delayed-recall decoder was swept across data from the MSR task, it revealed that the representation of stimulus category in all ROIs was priority dependent, and with a time course that was tightly coupled to the structure of the task: MVPA evidence for both categories was comparable at the beginning of the trial, prior to item prioritization, and closely tracked prioritization once prioritization cuing began. Coupled with the weak and uneven evidence for stimulus-level representation in frontal and parietal ROIs, these results are consistent with the idea that frontal and parietal networks were preferentially involved in controlling the maintenance of and changing of the priority of VOT-supported stimulus representations. Finally, the post hoc comparison of these MVPA time courses with the time courses of fluctuations in the baseline parameter of IEM suggest that the latter, too, may provide an index of the control of representations in visual WM.

### Dropping information from WM

Results from the final delay period of the MSR task indicated that IEMs of the DMI, in both conditions, yielded nonsignificant reconstructions with numerically negative amplitudes. Prompted by these trends, we undertook an exploratory analysis of the effect of the initial *relevance cue* on the neural representation of direction, because these retrocues also prompted the dropping of information from WM -- of a stimulus dimension (on *Unbound* trials); or of an entire stimulus (on *Bound* trials). On *Unbound* trials, on trials when the *relevance cue* indicated that “<Color>” was to be tested, the reconstruction of the two directions of motion returned to baseline levels (*p*s > .27; two-tailed; Figure 13B). However, on trials when the *relevance cue* indicated “<First>” or “<Second>” (i.e., *Bound* trials), the reconstruction of the direction of motion of the uncued (and, therefore, dropped) stimulus was significantly complementary to the trained model by the end of the ensuing delay (*p* < .01; two-tailed; Figure 13C).

**Figure 13.**
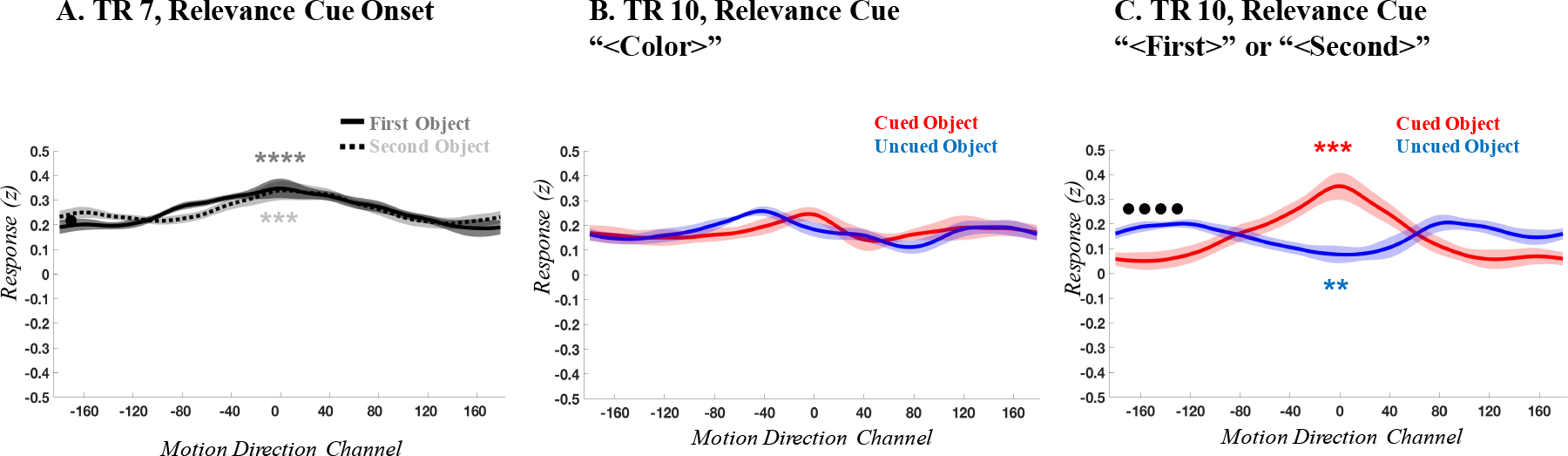
The effects of *relevance cued* “dropping” of direction information from WM. (A) Reconstructions of direction at TR 7, the time of onset of the *relevance cue*. (B) The reconstruction of direction, at TR 10, on trials when the *relevance cue* had specified “<Color>”, and so the direction associated with both objects could be dropped. (C) The reconstruction of direction, at TR 10, on trials when the *relevance cue* had specified “<First>” or “<Second>,” as a function of *relevance cued* status (i.e., belonging to the cued object (red), or belonging to the uncued object (blue)). All display conventions are the same as Figure 8.

## Discussion

The results from our study of a multiple serial retrocuing (MSR) task yielded three novel sets of empirical observations, each with important implications for our understanding of the mechanisms underlying visual working memory. The first relates to the effects of *prioritization cue 1* on stimulus representations. When two pieces of remembered information are drawn from separate objects, IEM indicates that the prioritization of one results in the strengthening of its neural representation, and in the weakening, to baseline levels, of the active neural representation of the unprioritized item (even though this UMI must be retained in working memory). When these two pieces of remembered information are drawn from the same object, in contrast, the effects of prioritization are markedly different: The active representation of the PMI is sustained, but does not increase in strength, and the strength of the active representation of the UMI declines, but nonetheless remains significantly above baseline. The second novel observation arises from contrasting the weak and uneven representation of stimulus identity in parietal and frontal cortex, whether assessed by IEM or MVPA, with the robust cue-locked dynamics of MVPA decoding of stimulus category in these two regions. The third set of novel observations relates to the removal of information from visual WM when it is no longer relevant for behavior, and provides preliminary evidence for an active “dropping” mechanism.

### Object-based attention in visual working memory

The differential pattern of results in the *Unbound* vs. the *Bound* conditions suggest that key principles governing object-based attentional prioritization in visual perception also apply to visual working memory. When the two remembered items belonged to separate objects (*Unbound*), the biasing of their competition for representation (Desimone & Duncan, 1995) resulted in the strengthening of the neural representation of the PMI at the expense of the strength of the UMI. When, however, the two remembered items belonged to the same object (*Bound*), the neural representation of the UMI remained elevated, consistent with an automatic spread of object-based attention to all elements of the remembered object. The theoretical implications of this finding are twofold. First, by illustrating object-based attention-like effects in visual working memory, they extend the boundary conditions for which it can be said that visual working memory appears to arise from “nothing more” than attention allocated to neural representations of objects not currently accessible to the eyes (c.f., Cowan, 1995; Chun, 2011; Myers, Stokes, & Nobre, 2017; Postle, 2006). Second, they support the idea that multidimensional objects are represented in visual working memory as bound objects, not as a collection of unbound features (e.g., Luck & Vogel, 1997; Luria & Vogel, 2011; Woodman & Vogel, 2008; Bays, Wu, & Husain, 2011; Wheeler & Treisman, 2002).

### Controlling priority in visual working memory

Stimulus representation in parietal and frontal cortex was markedly weaker than in the VOT ROI, whether assessed by IEM or by MVPA. There was, however, clear evidence that these two regions tracked the higher-order information of which stimulus category was prioritized during each epoch of the trial, and they did so with a high degree of temporal precision. This is consistent with the idea that cue-driven changes in priority were implemented in working memory via activity in frontoparietal circuits, whose representation of the prioritized category may have acted as a source of top-down bias on high-fidelity representations of stimulus features in VOT cortex.

### Evidence for the active removal of information from working memory

There are, in principle, two ways that information can exit working memory: its links to trial-specific context can decay after attention has been shifted away from it, or it can be actively removed, via suppression, recoding, or some other mechanism. Our results provide suggestive evidence for an active mechanism: In three instances when a retrocue indicated that the uncued information was no longer relevant for behavior, its neural representation transitioned from a robust “positive” reconstruction to one that approached (after *priority cue 2*) or achieved (after the *relevance cue*) a state that was “complementary” to the IEM of that item when it had been in the focus of attention. Although we cannot know from these data what mechanism(s) would have effected this change in representation (e.g., inhibition, recoding), we can postulate that they may only be engaged when task demands require the active removal of information from WM. In the 1-item delayed-recall task, in contrast, the neural representation of the sample item seems to just “fade away” at the end of the trial.

A mechanistically noncommittal interpretation of the retrocue-triggered transformation of DMIs is to suggest that once the identity of the information that will be relevant for the remainder of the MSR trial is known, the resultant DMI is processed in a manner that makes it least likely that it will interfere with performance on the remainder of the trial. A similar phenomenon has been observed in a different variant of the MSR task (with fMRI, Yu & Postle, 2018), in a 2-back task (with EEG, Wan, Cai & Postle, 2018) and in a dual serial visual search task (van Loon et al., 2018). It could be that recoding unprioritized information is a general mechanism for preventing that information from interfering with performance that needs to be guided by a PMI. By this account, removing an item from working memory would be accomplished in a two-step process: First, recode it so that it is less likely to interfere with the PMI; second, let the recoded representation decay.

## Author contributions

A.D.S., M.I.S., and B.R.P. designed research; A.D.S. and M.I.S performed research; A.D.S., M.I.S. analyzed data; and A.D.S, M.I.S. and B.R.P. wrote the paper.

## Supplementary Figures

**Supplementary Figure 1.**
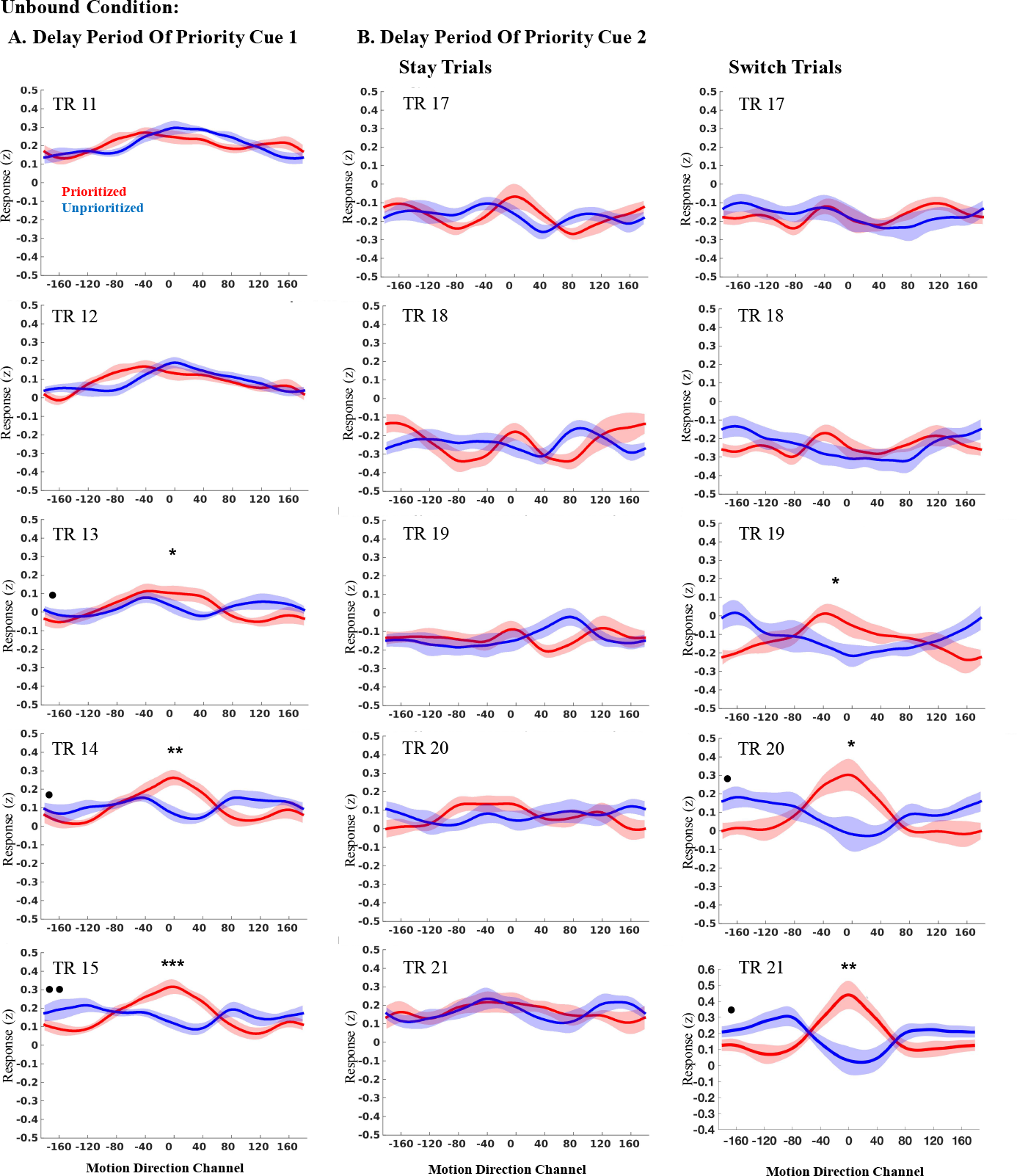
Individual time point plots for unbound trials in VOT of delay period of the priority cue 1. (A). Channel response is plotted on the Y-axis for individual time points, and the specific motion direction channel is plotted on the x-axis, centered around 0°. Each row is a set of reconstructions from a single time point starting with the priority cue 1 onset (TR 11) and ending with the probe onset (TR 15). Red traces represent reconstructions of directions on trials when direction was cued (prioritized reconstructions) and blue traces represent reconstructions of direction on trials when direction was uncued (unprioritized reconstructions). *Individual time point plots for unbound “stay” and “switch” trials in VOT of delay period of the priority cue 2* (B). Each row is a set of reconstructions from a single time point starting with the priority cue 2 onset (TR 17) and ending with the probe onset (TR 21). Significant differences between the prioritized and unprioritized motion directions for the amplitude (one-tailed bootstrap tested) and baseline (two-tailed bootstrap tested) estimates are indicated by the asterisks and dots respectively (* = *p*<.05: ** = *p* <.01; *** = *p* <.001; **** = *p* <.0001).

**Supplementary Figure 2.**
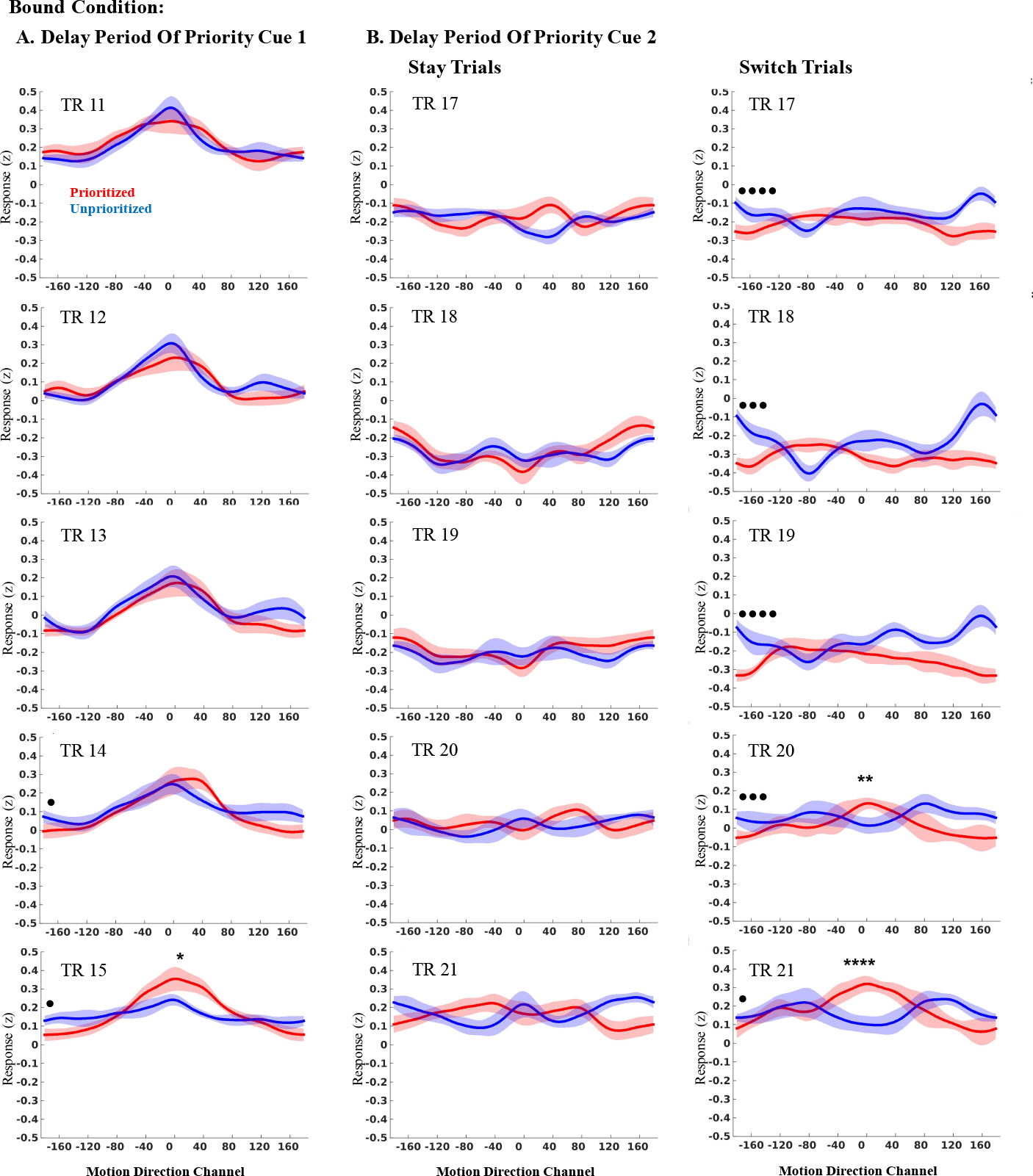
Individual time point plots for bound trials in VOT of delay period of the priority cue 1. (A) and *priority cue 2* (B). Conventions are the same as in Supplementary Figure 1.

## Footnotes

1 We use scare quotes when referring to a putative “dropping” operation, because our interpretations are agnostic with regard to whether no-longer-relevant information is actively removed from WM, or whether it passively decays out of WM.

## References

Bays, P. M., Catalao, R. F. G., & Husain, M. (2009). The precision of visual working memory is set by allocation of a shared resource. Journal of Vision, 9(10), 7.1–11. https://doi.org/10.1167/9.10.7

Bays, P. M., Wu, E. Y., & Husain, M. (2011). Storage and binding of object features in visual working memory. Neuropsychologia, 49(6), 1622–31. https://doi.org/10.1016/j.neuropsychologia.2010.12.023

Brincat, S. L., Siegel, M., Nicolai, C. Von, & Miller, E. K. (2017). Gradual progression from sensory to task-related processing in cerebral cortex.

Brouwer, G. J., & Heeger, D. J. (2009). Decoding and Reconstructing Color from Responses in Human Visual Cortex. Journal of Neuroscience, 29(44), 13992–14003. https://doi.org/10.1523/JNEUROSCI.3577-09.2009

Carandini, M., & Heeger, D. J. (2012). Normalization as a canonical neural computation. Nature Reviews Neuroscience, 13(1), 51.

Christophel, T. B., Iamshchinina, P., Yan, C., Allefeld, C., & Haynes, J.-D. (2018). Cortical specialization for attended versus unattended working memory. Nature Neuroscience, 21(April). https://doi.org/10.1038/s41593-018-0094-4

Chun, M. M. (2011). Visual working memory as visual attention sustained internally over time. Neuropsychologia, 49(6), 1407–1409. https://doi.org/10.1016/j.neuropsychologia.2011.01.029

Cox, R. W. (1996). AFNI : Software for Analysis and Visualization of Functional Magnetic Resonance Neuroimages, 173(29), 162–173.

D’Esposito, M., & Postle, B. R. (2015). The Cognitive Neuroscience of Working Memory. Annual Review of Psychology, 66(1), 115–142. https://doi.org/10.1146/annurev-psych-010814-015031

Desimone, R., & Duncan, J. (1995). Neural mechanisms of selective visual attention. Annual Review of Neuroscience, 18, 193–222. https://doi.org/10.1146/annurev.ne.18.030195.001205

Driver, J. (2001). A selective review of selective attention research from the past century. British Journal of Psychology, 92(1), 53–78. https://doi.org/10.1348/000712601162103

Duijnhouwer, J., Noest, A. J., Lankheet, M. J. M., Berg, A. V. Van Den, & Wezel, R. J. A. Van. (2013). Speed and direction response profiles of neurons in macaque MT and MST show modest constraint line tuning, 7(April), 1–11. https://doi.org/10.3389/fnbeh.2013.00022

Duncan, J. (1984). Selective attention and the organization of visual information. Journal of Experimental Psychology. General, 113(4), 501–17. https://doi.org/10.1037/0096-3445.113.4.501

Egly, R., Driver, J., & Rafal, R. D. (1994). Shifting visual attention between objects and locations: Evidence from normal and parietal lesion subjects. Journal of Experimental Psychology: General, 123(2), 161–177. https://doi.org/10.1037/0096-3445.123.2.161

Emrich, S. M., Riggall, A. C., Larocque, J. J., & Postle, B. R. (2013). Distributed Patterns of Activity in Sensory Cortex Reflect the Memory. Journal of Neuroscience, 33(15), 6516–6523. https://doi.org/10.1523/JNEUROSCI.5732-12.2013

Ester, E. F., Sprague, T. C., & Serences, J. T. (2015). Parietal and Frontal Cortex Encode Stimulus-Specific Mnemonic Representations during Visual Working Memory. Neuron, 87(4), 893–905. https://doi.org/10.1016/j.neuron.2015.07.013

Gazzaley, A., & Nobre, A. C. (2012). Top-down modulation: bridging selective attention and working memory. Trends in Cognitive Sciences, 16(2), 129–35. https://doi.org/10.1016/j.tics.2011.11.014

Griffin, I. C., & Nobre, A. C. (2003). Orienting attention to locations in internal representations. Journal of Cognitive Neuroscience, 15(8), 1176–94. https://doi.org/10.1162/089892903322598139

Kiyonaga, A., & Egner, T. (2013). Working memory as internal attention: toward an integrative account of internal and external selection processes. Psychonomic Bulletin & Review, 20(2), 228–42. https://doi.org/10.3758/s13423-012-0359-y

LaRocque, J. J., Lewis-Peacock, J. A., Drysdale, A. T., Oberauer, K., & Postle, B. R. (2013). Decoding Attended Information in Short-term Memory: An EEG Study. Journal of Cognitive Neuroscience, 25(1), 127–142. https://doi.org/10.1162/jocn_a_00305

Larocque, J. J., Lewis-Peacock, J. A., & Postle, B. R. (2014). Multiple neural states of representation in short-term memory? It’s a matter of attention. Frontiers in Human Neuroscience, 8(January), 1–14. https://doi.org/10.3389/fnhum.2014.00005

Larocque, J. J., Riggall, A. C., Emrich, S. M., & Postle, B. R. (2017). Within-Category Decoding of Information in Different Attentional States in Short-Term Memory, (October 2016), 4881–4890. https://doi.org/10.1093/cercor/bhw283

Lepsien, J., & Nobre, A. C. (2007). Attentional modulation of object representations in working memory. Cerebral Cortex, 17(9), 2072–2083. https://doi.org/10.1093/cercor/bhl116

Lewis-Peacock, J. A., & Postle, B. R. (2012). Decoding the internal focus of attention. Neuropsychologia, 50(4), 470–478. https://doi.org/10.1016/j.neuropsychologia.2011.11.006

Lewis-Peacock, J. A., Drysdale, A. T., Oberauer, K., & Postle, B. R. (2012). Neural evidence for a distinction between short-term memory and the focus of attention. Journal of Cognitive Neuroscience, 24(1), 61–79. https://doi.org/10.1162/jocn_a_00140

Lewis-Peacock, J. A., Drysdale, A. T., & Postle, B. R. (2014). Neural evidence for the flexible control of mental representations. Cerebral Cortex, 25(10), 3303–3313.

Luck, S. J., & Vogel, E. (1997). The capacity of visual working memory for features and conjunctions. Nature, 390(6657), 279–281. https://doi.org/10.1038/36846

Luria, R., & Vogel, E. K. (2011). Shape and color conjunction stimuli are represented as bound objects in visual working memory. Neuropsychologia, 49(6), 1632–1639. https://doi.org/10.1016/j.neuropsychologia.2010.11.031

Myers, N. E., Stokes, M. G., & Nobre, A. C. (2017). Prioritizing Information during Working Memory: Beyond Sustained Internal Attention. Trends in Cognitive Sciences, 21(6), 449–461. https://doi.org/10.1016/j.tics.2017.03.010

Nelissen, N., Stokes, M., Nobre, A. C., & Rushworth, M. F. S. (2013). Frontal and Parietal Cortical Interactions with Distributed Visual Representations during Selective Attention and Action Selection, 33(42), 16443–16458. https://doi.org/10.1523/JNEUROSCI.2625-13.2013

Oberauer, K., & Hein, L. (2012). Attention to Information in Working Memory. Current Directions in Psychological Science, 21(3), 164–169. https://doi.org/10.1177/0963721412444727

Pertzov, Y., Bays, P. M., Joseph, S., & Husain, M. (2013). Rapid forgetting prevented by retrospective attention cues. Journal of Experimental Psychology: Human Perception and Performance, 39(5), 1224–31. https://doi.org/10.1037/a0030947

Polyn, S. M., Natu, V. S., Cohen, J. D., & Norman, K. a. (2005). Category-specific cortical activity precedes retrieval during memory search. Science (New York, N.Y.), 310(5756), 1963–6. https://doi.org/10.1126/science.1117645

Pribram KH, Ahumada A, Hartog J, Roos L (1964). A progress report on the neurological processes disturbed by frontal lesions in primates. In: The frontal granular cortex and behavior (Warren JM, Akert K, eds), pp 28–55. New York: McGraw-Hill Book Company

Rose, N. S., LaRocque, J. J., Riggall, A. C., Gosseries, O., Starrett, M. J., Meyering, E. E., & Postle, B. R. (2016). Reactivation of latent working memories with transcranial magnetic stimulation. Science, 354(6316), 1136–1139.

Sahan, M. I., Verguts, T., Boehler, C. N., Pourtois, G., & Fias, W. (2016). Paying attention to working memory: Similarities in the spatial distribution of attention in mental and physical space. Psychonomic bulletin & review, 23(4), 1190–1197.

Serences, J. T., & Saproo, S. (2012). Neuropsychologia Computational advances towards linking BOLD and behavior. Neuropsychologia, 50(4), 435–446. https://doi.org/10.1016/j.neuropsychologia.2011.07.013

Sheldon, A.D., Saad, E., Sahan, M. I., Meyering, E., Postle, B. R. (2017). Regional Heterogeneity of the Effects of Attentional Prioritization. Cogntivie Neuroscience Society, March 2017, San Francisco, CA.

van Loon, A. M., Olmos-solis, K., Fahrenfort, J. J., & Olivers, N. C. (2018). Current and future goals are represented in opposite patterns in object-selective cortex, 1–25.

Vecera, S. P., Behrmann, M., & McGoldrick, J. (2000). Selective attention to the parts of an object. Psychonomic Bulletin & Review, 7(2), 301–308. https://doi.org/10.3758/BF03212985

Wan, Q, Cai, Y., and Postle, B. R. (2018). Tracking stimulus representation across a 2-back visual working memory task. Society for Neuroscience, November 2018, San Diego, CA.

Wang, X. H. X., Merriam, X. E. P., Freeman, J., & Heeger, X. D. J. (2014). Motion Direction Biases and Decoding in Human Visual Cortex, 34(37), 12601–12615. https://doi.org/10.1523/JNEUROSCI.1034-14.2014

Wheeler, M. E., & Treisman, A. M. (2002). Binding in short-term visual memory. Journal of Experimental Psychology: General, 131(1), 48–64. https://doi.org/10.1037//0096-3445.131.1.48

Woodman, G. F., & Vogel, E. K. (2008). Selective storage and maintenance of an object’s features in visual working memory. Psychonomic Bulletin & Review, 15(1), 223–229. https://doi.org/10.3758/PBR.15.1.223

Yu, Q., & Postle, B. R. Different states of priority recruit different neural codes in visual working memory. bioRxiv, doi:https://doi.org/10.1101/334920

Zhang, Y., Meyers, E. M., Bichot, N. P., Serre, T., Poggio, T. A., & Desimone, R. (2011). Object decoding with attention in inferior temporal cortex, 108(21), 8850–8855. https://doi.org/10.1073/pnas.1100999108

Zokaei, N., Manohar, S., Husain, M., & Feredoes, E. (2013). Causal Evidence for a Privileged Working Memory State in Early Visual Cortex. Journal of Neuroscience, 34(1), 158–162. https://doi.org/10.1523/JNEUROSCI.2899-13.2014

